# Structural and Energetic Determinants of Monobody Recognition of Oncogenic KRAS Variants

**DOI:** 10.64898/2026.07.09.737552

**Authors:** Amit Kumar, Yu-ming M. Huang

## Abstract

Monobodies are engineered binding proteins that recognize extended protein surfaces and offer advantages over small-molecule inhibitors for targeting challenging KRAS oncoproteins. Monobody 12D4 exhibits high affinity and selectivity for the oncogenic KRAS(G12D) mutant, but the molecular determinants governing its recognition and the basis for its mutant selectivity remain poorly understood. Here, we combined molecular dynamics simulations and energy calculations to characterize the interactions between monobody 12D4 and WT KRAS as well as four clinically relevant oncogenic variants (G12C, G12D, G12V, and G12R) in both GTP-and GDP-bound states. Our simulations revealed that 12D4 recognition depends on a conserved hydrophobic interaction network centered on the monobody FG loop (residues L77, F78, and W79). This network forms stable contacts with KRAS Switch II and α3-helix. The energy calculations also showed that residue K75 of 12D4 formed a mutation-specific electrostatic interaction with KRAS G12D. This interaction contributed significantly to the affinity of 12D4 toward this mutant, whereas this interaction was absent in other variants. No monobody currently exists for targeting KRAS G12R in either nucleotide state, and no monobody selectively targets KRAS G12C and G12V in the GDP-bound inactive state. To address these, we performed computational redesign at residues 75. We identified mutations (K75Q, K75Y, and K75M) that enhanced predicted binding to G12C, G12R, and G12V variants through reorganization of interfacial contacts. Our work establishes a structural framework for understanding KRAS-monobody recognition and provides a rational foundation for engineering variant-selective monobodies with improved affinity toward previously untargetable KRAS mutants.

## Introduction

Monobodies are engineered binding proteins derived from the human fibronectin type III (FN3) scaffold, a small and stable domain structurally analogous to antibody variable regions but lacking disulfide bonds^1^. Their thermal stability and single-domain architecture enable engineering for high affinity and specificity using directed evolution strategies, such as yeast or phage display^2^. Unlike small-molecule inhibitors, which typically bind in a well-defined hydrophobic pocket, monobodies recognize extended and dynamic protein surfaces, including protein-protein interaction (PPI) interfaces^3^. This feature allows them to target epitopes that are generally inaccessible to conventional small ligands, which are prone to limited selectivity, off-target toxicity, and resistance^4,5^. The modularity of the FN3 scaffold further permits the development of mutation-specific binders, engage cryptic allosteric sites, and design of bispecific or degrader constructs, thereby broadening the repertoire of therapeutically tractable targets in oncology^4,6–8^.

KRAS encodes a small GTPase that functions as a molecular switch in the RAS/MAPK signaling pathway^9^. In addition, KRAS regulates the RAS/MAPK pathway, the PI3K/AKT/mTOR pathway (promoting cell survival and metabolism), the RAL-GDS/RAL pathway (driving invasion and metastasis), and the TIAM1-RAC1 pathway (modifying cell shape and adhesion)^10–12^. The alternating cycles between active GTP-bound and inactive GDP-bound states govern cellular processes, such as proliferation, differentiation, and apoptosis. Structurally, KRAS contains several key functional regions, including the phosphate-binding loop (P-loop, residues 10-17), Switch I (residues 30-40), and Switch II (residues 56-78) (**Figure 1**), which undergo conformational changes upon nucleotide exchange and mediate effector binding^9,13,14^. Extensive computational and experimental studies have characterized the dynamics of these regions and their roles in KRAS regulation^9,13–16^.

**Figure 1.**
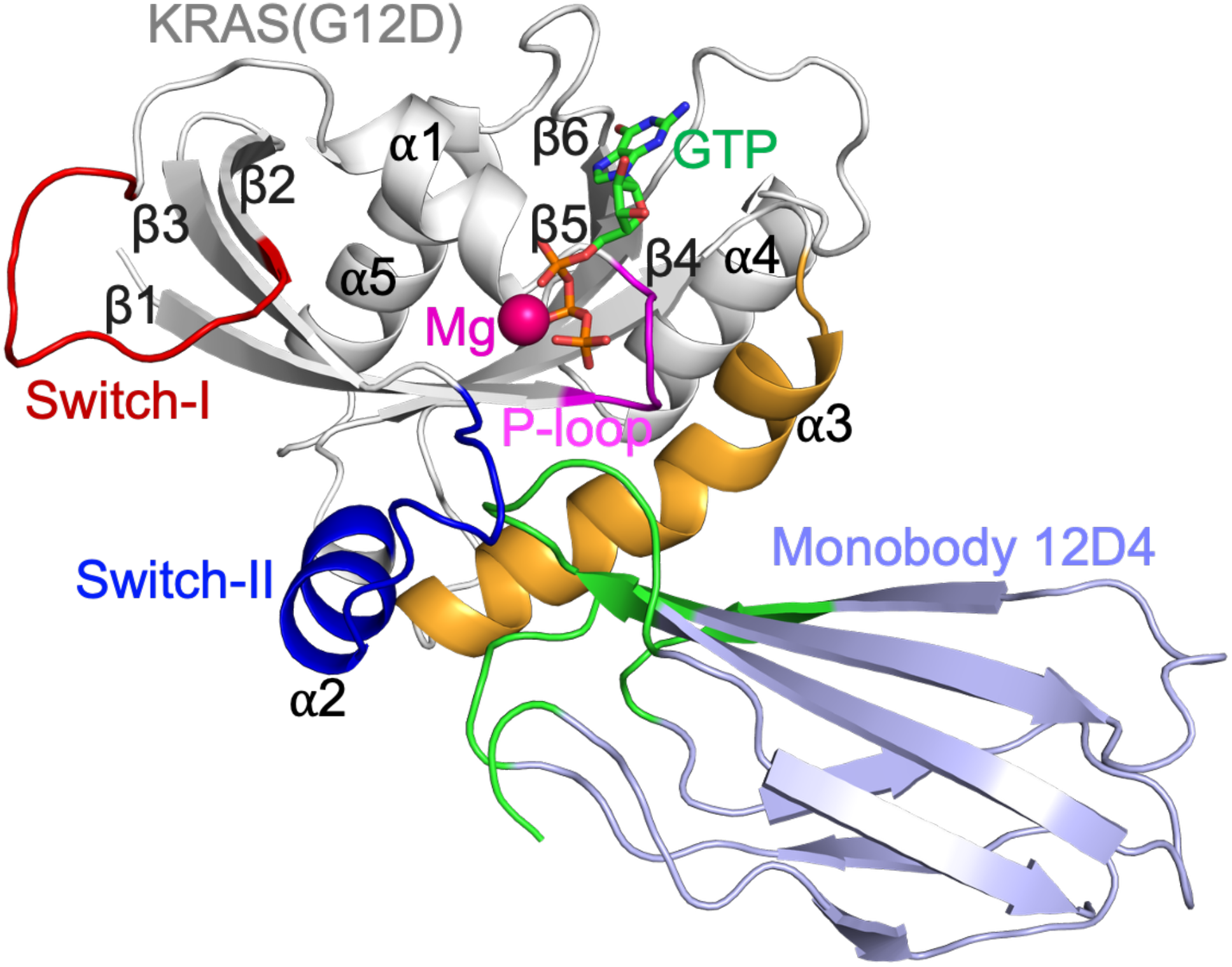
Modeled structure of the KRAS(G12D)(GTP) variant (gray) in complex with the monobody 12D4 (purple). Key structural elements are highlighted: P-loop (residues 10-14) in magenta, Switch I (residues 30-40) in red, Switch II (residues 58-72) in blue, and α3-helix (residues 87-104) in orange. Monobody residues involved in interface interactions are shown in green.

Oncogenic mutations, particularly at codon 12 (G12D, G12C, G12V, and G12R), impair both intrinsic and GAP-stimulated GTP hydrolysis, with reduced GAP-mediated hydrolysis contributing substantially to the persistent activation of KRAS^17,18^. Recent studies showed that the G12C mutation alters the conformational dynamics of the Switch I and Switch II regions of KRAS^19–21^. These mutants lock KRAS in its active state and lead to persistent activation of downstream effectors, notably the RAF-MEK-ERK and PI3K-AKT cascades^18,22^. The G12D mutation, prevalent in pancreatic, colorectal, and lung cancers, introduces a negatively charged residue into the P-loop, which stabilizes the active KRAS conformation then promotes oncogenic signaling and therapy resistance^23–26^. Although covalent KRAS(G12C) inhibitors, such as sotorasib and adagrasib^27^, have achieved clinical approval, their sustained efficacy can be limited by the emergence of acquired resistance during treatment^28^. In addition, other KRAS mutants remain largely untargetable due to their smooth surfaces and high GTP affinity^22^ (**Figure 1**). Noncovalent inhibitors, including MRTX1133 and ASP3082, show promise but still face challenges in potency and mutant selectivity^29,30^. In contrast, macromolecular binders, e.g., nanobodies and monobodies, can interact with extended epitope surfaces with high specificity^4,31^.

Several monobodies have recently been developed to selectively target oncogenic KRAS variants^32,33^. Among them, 12D4 exhibits the highest reported affinity for KRAS(G12D), with 400-fold greater affinity than WT KRAS^32,33^, making it one of the most promising scaffolds for variant-selective targeting. However, its limited solubility has prevented experimental structure determination. As a result, structural information is currently limited to the closely related monobody variants 12D1 and 12D5 in complex with KRAS(G12D)^32^. Recent computational analysis further showed that 12D1 and 12D5 can restore altered KRAS(G12D) conformational transitions and allosteric communication pathways^34^. These structures reveal that monobody recognition is mediated primarily through extensive electrostatic and hydrophobic interactions with the Switch II and nearby helical regions^32^. Whether 12D4 adopts a similar binding mode or uses distinct molecular interactions to achieve its superior affinity remains unknown. Furthermore, the structural basis of its recognition of other clinically relevant KRAS variants, including G12C, G12V, and G12R, has not been established.

Monobodies 12D4 and 12D5 are closely related variants derived from the FN3 scaffold and differ by only two sequence features. At position 51, 12D4 contains a Ser residue, whereas 12D5 carries a Pro substitution. In addition, 12D5 includes a short C-terminal EIDK extension that is absent in 12D4^32^. Despite these minimal sequence differences, the two monobodies exhibit distinct binding behavior toward KRAS(G12D). In particular, 12D4 binds KRAS(G12D) with substantially higher affinity, while 12D5 displays a 4-fold reduction in binding strength^32^. This observation suggests that the identity of residue 51 and the native C-terminal context can influence productive target engagement, either by directly contributing to the binding interface or by subtly altering the structural presentation of the binding loops. Because of its superior affinity and selectivity for KRAS(G12D), 12D4 was selected for further investigation to understand how these subtle sequence features enhance recognition of the mutant KRAS protein.

In this study, we constructed structural models of 12D4 in complex with WT KRAS and four oncogenic variants (G12D, G12C, G12V, and G12R) and investigated their interactions using Gaussian accelerated molecular dynamics (GaMD) simulations^35^. The simulations were combined with MM/PBSA calculations^36^, residue-level interaction analyses, and thermodynamic integration (TI)^37^ to characterize the conformational and energetic determinants of monobody recognition. Beyond defining the molecular basis for the exceptional affinity of 12D4 toward KRAS(G12D), we sought to identify sequence features that could be exploited to improve binding to other KRAS variants for which selective monobody binders remain unavailable or underdeveloped. Using computational protein design guided by TI calculations, we further evaluated mutations predicted to enhance recognition of KRAS(G12C), KRAS(G12V), and KRAS(G12R). Our studies provide a structural and energetic framework for the rational development of variant-selective monobodies targeting oncogenic KRAS.

## Methods

### Molecular Models

The initial structure of the KRAS(G12D)-12D4 complex was constructed using the crystal structure of the KRAS(G12D)-12D5 complex (PDB ID: 8F0M) as the template. Chain C (KRAS(G12D)-GTPγS) and chain D (monobody 12D5) were used as the starting structural framework. P51 in 12D5 was replaced with S51, and the C-terminal EIDK extension present in 12D5 but absent in 12D4 was removed^32^. Missing residues were reconstructed using MODELLER^38^. The GDP-bound model was subsequently generated from the same GTPγS)-bound structural framework, such that GTP-and GDP-bound simulations shared the same initial protein-monobody conformation. The resulting model was used for energy minimization prior to simulation. Structures of WT KRAS and the G12C, G12V, and G12R variants were then generated by introducing the corresponding point mutations into the KRAS(G12D)-12D4 model.

### Molecular Dynamics Simulation Protocol

All simulations were performed using the AMBER20 simulation package^39^. The protein was described using the ff19SB force field^40^. Parameters for GTP and GDP were generated using the General Amber Force Field (GAFF)^41^. Na^+^ and Cl^−^ ions were modeled using the Joung-Cheatham ion parameters^42^. The Mg²⁺ ion was modeled using the previously published Li-Merz parameters^43^, which were directly adopted from the force field implemented in AMBER20. The initial minimization consisted of 500 steps applied to hydrogen atoms, followed by 5000 steps applied to hydrogen atoms and side chains, and an additional 5000 steps applied to the entire protein. Each KRAS-12D4 complex was then solvated in an explicit TIP3P^44^ water box with a minimum distance of 12 Å between the protein and the box boundary. Counterions were added to neutralize the system, and additional Na^+^ and Cl^−^ ions were introduced to obtain a physiological salt concentration of 150 mM. After solvation, a second minimization was performed using 1000 steps for the solvent molecules followed by 5000 steps for the complete system. The minimized systems were gradually heated from 50 to 300 K in 50 K increments under the NVT ensemble. Each intermediate temperature was maintained for 10 ps, followed by 100 ps of equilibration at 300 K. Conventional MD simulations were then carried out for 20 ns under isothermal-isobaric (NPT) conditions at 300 K and 1 atm to ensure stable equilibration before enhanced sampling. Temperature was regulated using a Langevin thermostat with a collision frequency of 5 ps^−1^ ^45^. Long-range electrostatic interactions were treated using the particle mesh Ewald (PME) method with a nonbonded cutoff of 12 Å^46^. Covalent bonds involving hydrogen atoms were constrained using the SHAKE algorithm^47^, which permitted a 2 fs integration time step.

### Gaussian Accelerated Molecular Dynamics Simulations

Gaussian accelerated molecular dynamics (GaMD) is an enhanced sampling method that accelerates conformational transitions by adding a harmonic boost potential to smooth the system potential energy surface^48,49^. The simulations were performed using the dual-boost scheme, in which boost potentials were applied to both the total potential energy and the dihedral potential energy. Each simulation consisted of a 52 ns preparation stage followed by a 1 μs production simulation. The preparation stage included a 2 ns conventional MD simulation to collect potential energy statistics, a 1 ns GaMD simulation with an initial boost potential, and a 49 ns GaMD equilibration during which the boost parameters were updated every 500 ps. The production simulations were then carried out for 1 μs using fixed boost parameters. The lower-bound threshold energy was used for boost calculation, and the upper limit of the boost potential standard deviation was set to 6.0 kcal mol⁻¹ for both the total and dihedral potential energies. All GaMD simulations were performed under NPT conditions using the same thermostat, barostat, electrostatic setting, and integration parameters as the conventional MD simulations. Trajectory snapshots were saved every 10 ps for subsequent analysis. Three independent GaMD replicas with different initial velocity distributions were performed for each system to improve statistical sampling.

### Post-MD Analysis

Visual Molecular Dynamics (VMD)^50^ and PyMOL^51^ were used to visualize the simulation systems and create figures. Trajectory analyses were primarily performed using the CPPTRAJ program^52^, including calculations of the root mean squared fluctuation (RMSF), root mean squared deviation (RMSD), and radius of gyration (Rg). RMSF was calculated using the alpha carbon (Cα) atom of each residue, whereas RMSD was calculated using the protein backbone atoms (C, CA, N, O). The percentage of contact time between KRAS residues and 12D4 was calculated using MDAnalysis^53^ with a distance cutoff of 3.5 Å. All reported values represent averages from three independent simulation replicas. For conformational analysis of the KRAS switch regions, the trajectories were clustered in CPPTRAJ^52^ using the center-of-mass distance between Switch I (residues 30-40) and Switch II (residues 56-78). The center-of-mass distance was calculated from three independent trajectories, with snapshots saved every 1 ns, and subsequently subjected to K-means clustering to partition the conformational ensemble into distinct macrostates.

### Relative Binding Free Energy Calculations

Relative binding free energies (ΔΔG) of monobody mutations were calculated using thermodynamic integration (TI) as implemented in AMBER20^37,39^. For each mutation, an alchemical transformation was performed for the monobody in both the KRAS-bound complex and the unbound monobody in solution using the thermodynamic cycle shown in **Figure S1**. The relative binding free energy was calculated as

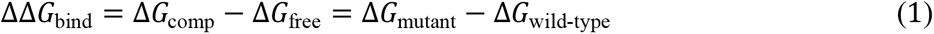

Where (Δ*G*_comp_) and (Δ*G*_free_) are the free energy changes associated with the alchemical transformation in the bound and unbound states, respectively. A one-step alchemical protocol was employed in which the electrostatic and van der Waals interactions of the transforming residue were changed simultaneously. Each λ window was energy minimized using 5000 steps of steepest descent followed by 5000 steps of conjugate gradient minimization. The system was then heated from 0 to 300 K in 50 K increments under the NVT ensemble, equilibrated for 1 ns under NPT conditions, and followed by a 5 ns production simulation for free energy collection. A total of 21 evenly spaced λ windows were used for each transformation. For each mutation, three independent free-energy calculation replicas were performed using these 21 λ windows. Convergence was assessed by comparing the free-energy estimates across the three replicas, with statistical uncertainties below 1 kcal/mol for all mutations. Free energy integration was performed using Gaussian quadrature. All mutations were carried out in the forward direction and repeated three times with independent initial velocity distributions. Reverse transformations were not performed because mutations involving increases in side-chain size generally require substantially longer sampling to achieve convergence and often exhibit significant hysteresis^54–57^. The reported ΔΔG values represent the mean of three independent calculations, and the corresponding errors are reported as the standard error of the mean.

### Protein-protein Interaction Energy Calculations

Binding energies between KRAS and 12D4 were calculated using the molecular mechanics Poisson-Boltzmann surface area (MM/PBSA) method implemented in AmberTools^36,58^. For each system, the last 500 ns of the GaMD trajectory were analyzed by extracting snapshots every 1 ns. The binding energy was calculated as the sum of the molecular mechanics interaction energy and the solvation free energy,

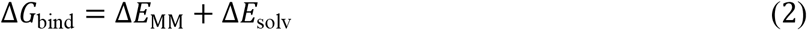

where the molecular mechanics term consists of electrostatic and van der Waals interactions, and the solvation free energy includes polar and nonpolar contributions. Polar solvation energies were calculated using the Poisson-Boltzmann model with dielectric constants of 4.0 for the solute and 80 for the solvent. A solvent probe radius of 1.4 Å was used throughout the calculations. Pairwise electrostatic and van der Waals interaction energies between interfacial residues of KRAS and the 12D4 were calculated using the NAMDEnergy plugin implemented in VMD^50^. All reported energies were calculated from three independent GaMD trajectories. Error bars represent the standard error of the mean.

## Results

### Structural Stability and Conformational Flexibility of KRAS-12D4 Complexes

We performed 1 μs all-atom GaMD simulations of WT and oncogenic KRAS variants (G12C, G12D, G12V, and G12R) in complex with monobody 12D4 in explicit solvent. Global stability of the complexes was evaluated by calculating backbone RMSD relative to the initial energy-minimized structures (**Figure S2**). Across all systems, the monobody remained structurally stable, with mean RMSD values below 2.23 Å (**Tables S1 and S2**). In the GTP-bound active state, monobody fluctuations ranged from 1.85 to 2.23 Å (**Table S1**), while the GDP-bound systems showed comparable stability with RMSD values between 1.65 and 2.14 Å (**Table S2**). The structural integrity of the monobody was further supported by radius of gyration (Rg) analysis, which remained stable with an average value of 13 Å throughout the simulations (**Figure S3**). This result indicates that the monobody maintains a compact β-sandwich fold and remains the intrinsic rigidity of the scaffold during complex formation regardless of the KRAS mutation or nucleotide state (GTP or GDP).

In contrast to the rigid monobody, KRAS exhibited greater conformational flexibility, particularly in the GTP-bound active state. Backbone RMSD analysis revealed noticeable variation among the mutants, which indicates mutation-dependent differences in structural dynamics. The RMSD for KRAS(GTP) varied significantly among mutants, e.g., RMSD of G12C and G12R is 4.42 and 2.70 Å, respectively (**Table S1**). In the GDP-bound inactive state, overall fluctuations were slightly reduced with RMSD ranging from 2.31 to 4.09 Å (**Table S2**). Despite these differences in RMSD, the Rg of KRAS remained stable at 15.5 Å across all complexes (**Figure S3**). This suggests that the global fold of the protein is preserved while conformational changes are primarily localized within functional loop regions.

To identify which regions of KRAS contribute to the structural fluctuations, we performed a subdomain RMSD analysis. The analysis focused on key regions near the protein-protein interface, including the P-loop, α3-helix, Switch II, and the highly mobile Switch I region (**Tables S1 and S2**). Among these elements, Switch I showed the largest RMSD values and emerged as the primary contributor to structural heterogeneity. This localized flexibility was further supported by residue-level RMSF analysis, which consistently identified Switch I as the most dynamic region of KRAS in both the GTP-bound (active) and GDP-bound (inactive) states across all variants (**Figure 2**). In contrast, the monobody exhibited high fluctuations primarily in its flexible loops, which are located away from the KRAS-binding interface (**Figure 2**).

**Figure 2.**
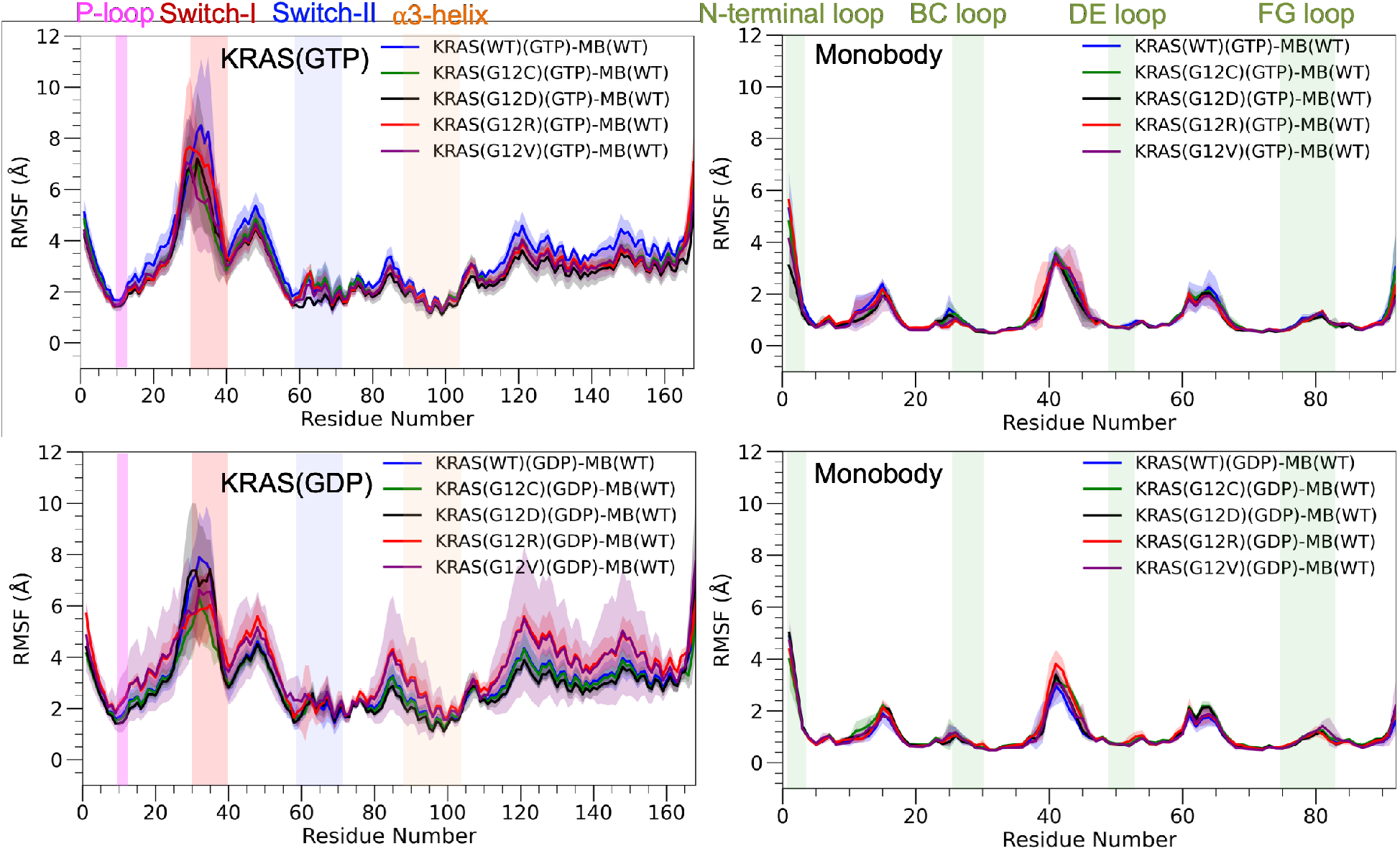
Root-mean-square fluctuation (RMSF) profiles of five KRAS system (WT, G12D, G12C, G12R, and G12V) in complex with the monobody 12D4, shown for both active (GTP-bound) and inactive (GDP-bound) states. Key KRAS structural elements are highlighted: P-loop (magenta), Switch I (red), Switch II (cyan), and α3-helix (orange). Monobody residues involved in interface interactions are highlighted in green.

### Conformational Changes in KRAS Switch Regions

The Switch I and Switch II regions play central roles in KRAS signaling. Residue-level fluctuation analysis identified Switch I as the most dynamic region in all systems (**Figure 2**). We therefore next examined the relative motions of the two switch regions to characterize mutation-and nucleotide-dependent conformational changes in KRAS by measuring the center-of-mass distance between Switch I and Switch II. This distance quantifies the relative positions of the two switch regions. Based on the distance distributions, we defined three conformational states: closed (<15 Å), open (15-20 Å), and wide-open (>20 Å) (**Figure 3A**). **Figure 3 and S4** show that all KRAS systems have multiple conformational states, although the relative populations differ among mutants and nucleotides.

**Figure 3.**
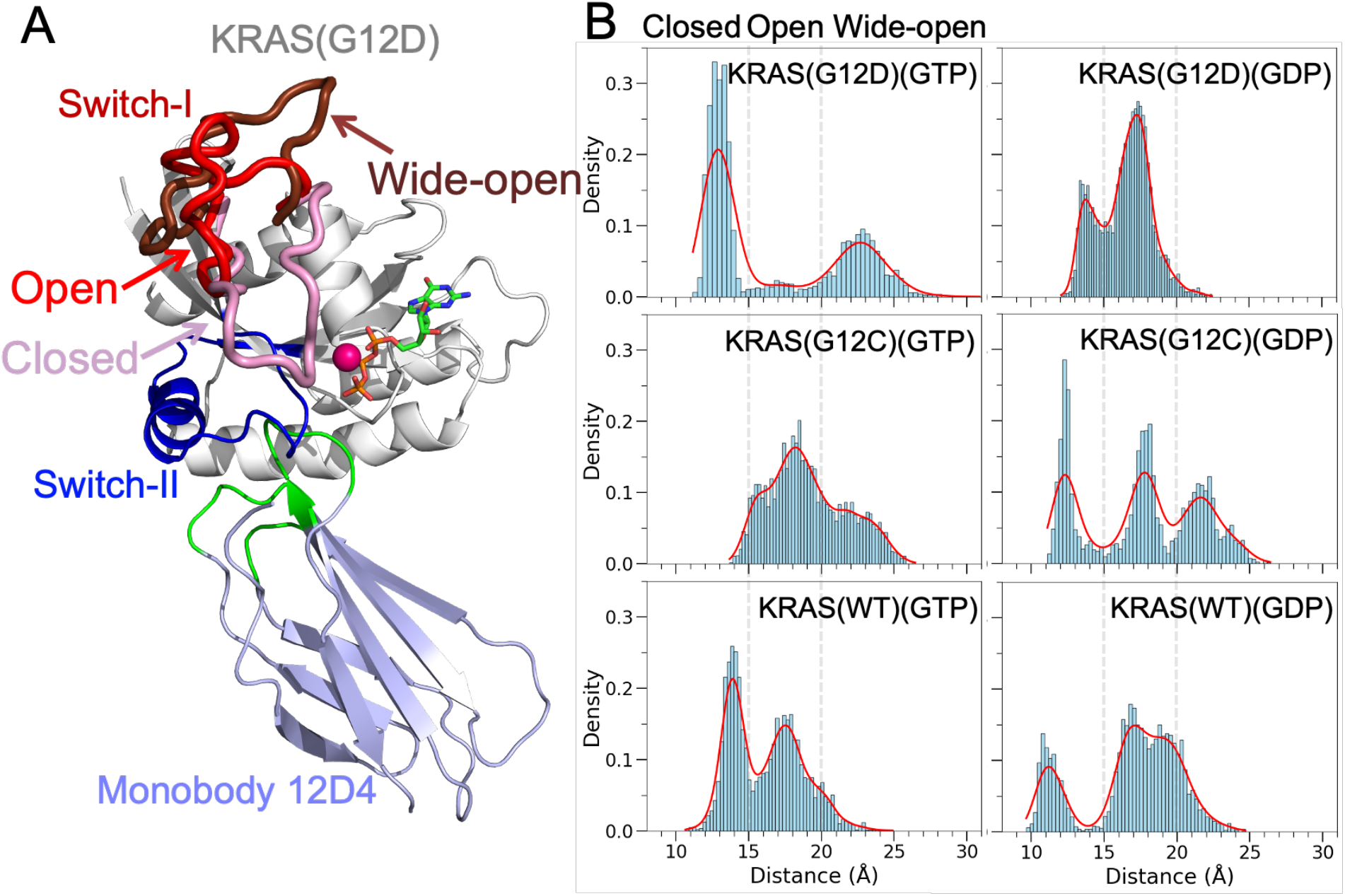
Conformational analysis of KRAS switch regions. (A) Representative conformations illustrating closed (pink), open (red), and wide-open (dark red) states of KRAS Switch I are highlighted. (B) Probability density distributions show the center-of-mass distances between Switch I and Switch II for KRAS G12D, G12C, and WT in GTP-and GDP-bound states.

In the GTP-bound systems, KRAS(G12D), KRAS(G12V), and KRAS(WT) preferentially populate the closed state, with the dominant population centered near 13-15 Å **(Figure 3B and S4)**. KRAS(G12C)(GTP) and KRAS(G12R)(GTP) exhibit a shift toward more open conformations, with the largest populations observed between 15-20 Å. The GDP-bound systems display a different conformational preference. Open conformations become the dominant in WT, G12D, G12R, and G12V. The populations of these systems occur near 16-18 Å. KRAS(G12C)(GDP) exhibits the broadest distribution and samples the closed, open, and wide-open states with similar populations. In the context of KRAS-monobody complexes, our results demonstrate that nucleotide state strongly influences switch-region organization. GTP binding favors more compact conformations, whereas GDP binding shifts the equilibrium toward more open arrangements. The G12C and G12R mutations further enhance switch-region flexibility, while G12D preferentially stabilizes compact conformations, particularly in the active GTP-bound state. These findings describe the conformational behavior of monobody-bound KRAS and should not be directly generalized to free KRAS or KRAS bound to small-molecule inhibitors, whose nucleotide and mutation dependent structural and dynamic properties have been extensively studied previously^59^.

### Binding Energies of KRAS-12D4 Complexes

To elucidate the thermodynamic basis of monobody binding to KRAS(WT) and G12 variants, we performed an analysis of binding free energies (*ΔG*_bind_) using the MMPBSA method^58^ for both GTP-and GDP-bound states. Overall, the 12D4 monobody exhibits the strongest binding affinity toward the G12D mutant (**Figure 4**). When comparing nucleotide states within the same variant, G12D and G12C show similar binding affinities in both GTP-and GDP-bound forms. In contrast, for G12R and G12V, the monobody binds more strongly to the GTP-bound (active) state. For KRAS(WT), binding is stronger in the GDP-bound (inactive) state.

**Figure 4.**
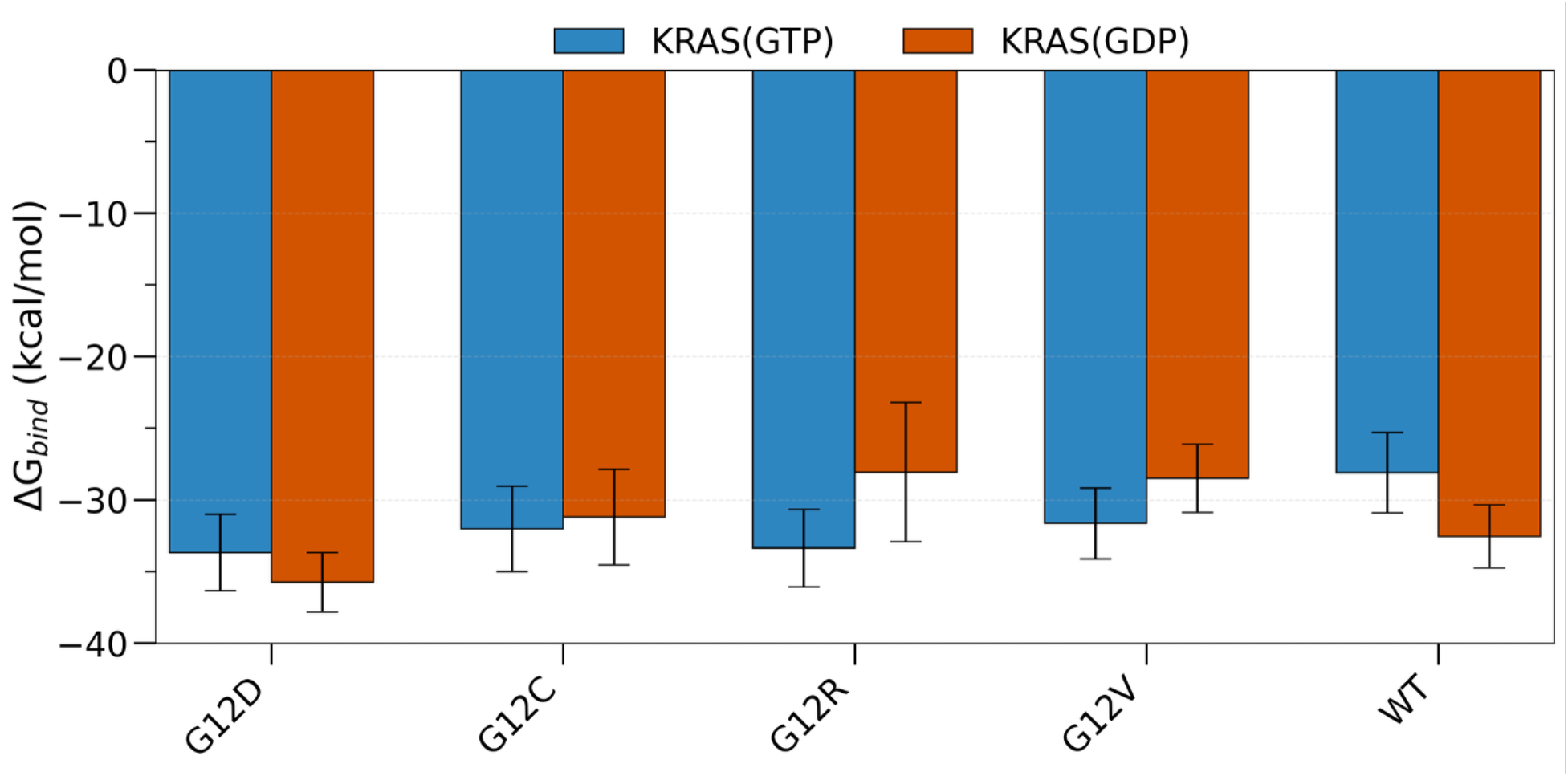
Binding free energy (Δ*G_bind_*) obtained from MM/PBSA calculations. The energy is computed for the final 500-ns simulations of KRAS-12D4 complexes in the GTP-bound active and GDP-bound inactive states. The error bars indicate the standard deviation calculated from three independent MD simulations.

In the GTP-bound state, all oncogenic variants exhibited a more favorable binding than the KRAS(WT)(GTP) (**Figure 4 and Table S3**). The KRAS(G12D) mutant formed the most stable complex, with a *ΔG*_bind_ value of -33.68 kcal/mol, which is 5.59 kcal/mol lower than the WT (-28.09 kcal/mol). The G12R and G12C variants also displayed moderate binding affinity, while the G12V mutation showed the weakest monobody binding affinity among the mutants. These results suggest that the monobody possesses a broad-spectrum efficacy for GTP-bound oncogenic KRAS, with a clear preference for the Asp and Arg substitutions at the 12^th^ position.

In the GDP-bound state, the binding affinity of KRAS and monobody changes, particularly for the G12R and G12V mutants. The KRAS(G12D)(GDP) complex still maintained the strongest binding affinity with the monobody among all systems, with an *ΔG*_bind_ of -35.75 kcal/mol (**Figure 4 and Table S4**). In contrast, the KRAS(G12R)(GDP) variant showed a significant decrease in binding affinity (-28.07 kcal/mol), which suggests that the Arg substitution destabilizes the GDP-bound state interface. Similarly to the GTP-bound state, KRAS(G12V)(GDP) showed weaker binding with the monobody. Our results indicate that the ability of the monobody to recognize the inactive state is highly sensitive to the specific side-chain of the 12^th^ residue, with Arg and Val mutation reducing interface stability.

### Residue-Specific Energetics and Contact Stability at the KRAS-12D4 Interface

To determine how individual residues of 12D4 contribute to binding affinity for KRAS, we performed residue-level interaction energy decomposition and contact occupancy analyses at the protein-protein interface for WT KRAS and its oncogenic variants in both GTP-and GDP-bound states. The interaction energy was decomposed into electrostatic and van der Waals components to quantify residue-specific contributions. Time-averaged contact occupancies were calculated to distinguish persistent contacts from transient interactions. These complementary analyses provide a quantitative description of how noncovalent interactions and contact stability define the binding interface. High-occupancy contacts represent key interaction hotspots that contribute to binding affinity.

Overall, 12D4 interacts with KRAS through four loop regions, including the N-terminal loop (residues 1-8), BC loop (residues 22-29), DE loop (residues 52-55), and FG loop (residues 77-85) (**Figures 1, 2, 5, and S5**). KRAS interacts with 12D4 primarily through the P-loop, Switch I, Switch II, and the α3-helix (**Figures 6 and S6**). Among these regions, Switch II and the α3-helix form the most stable contacts and constitute the core of the binding interface, as indicated by their high contact occupancies. Interaction energy analysis supports this observation. The strongest contributions localize to the 12D4 N-terminal loop (V1) and FG loop (F78, W79, S80) (**Figures 5 and S5**). V1 and S80 mainly contribute electrostatic interactions, whereas F78 and W79 provide dominant van der Waals stabilization. The FG loop serves as the primary structural determinant of binding. It maintains high-occupancy contacts with the Switch II region across all KRAS variants. In addition, the BC loop, particularly residues V27 and T28, provides secondary stabilization through combined electrostatic and van der Waals interactions (**Figures 5 and S5**).

**Figure 5.**
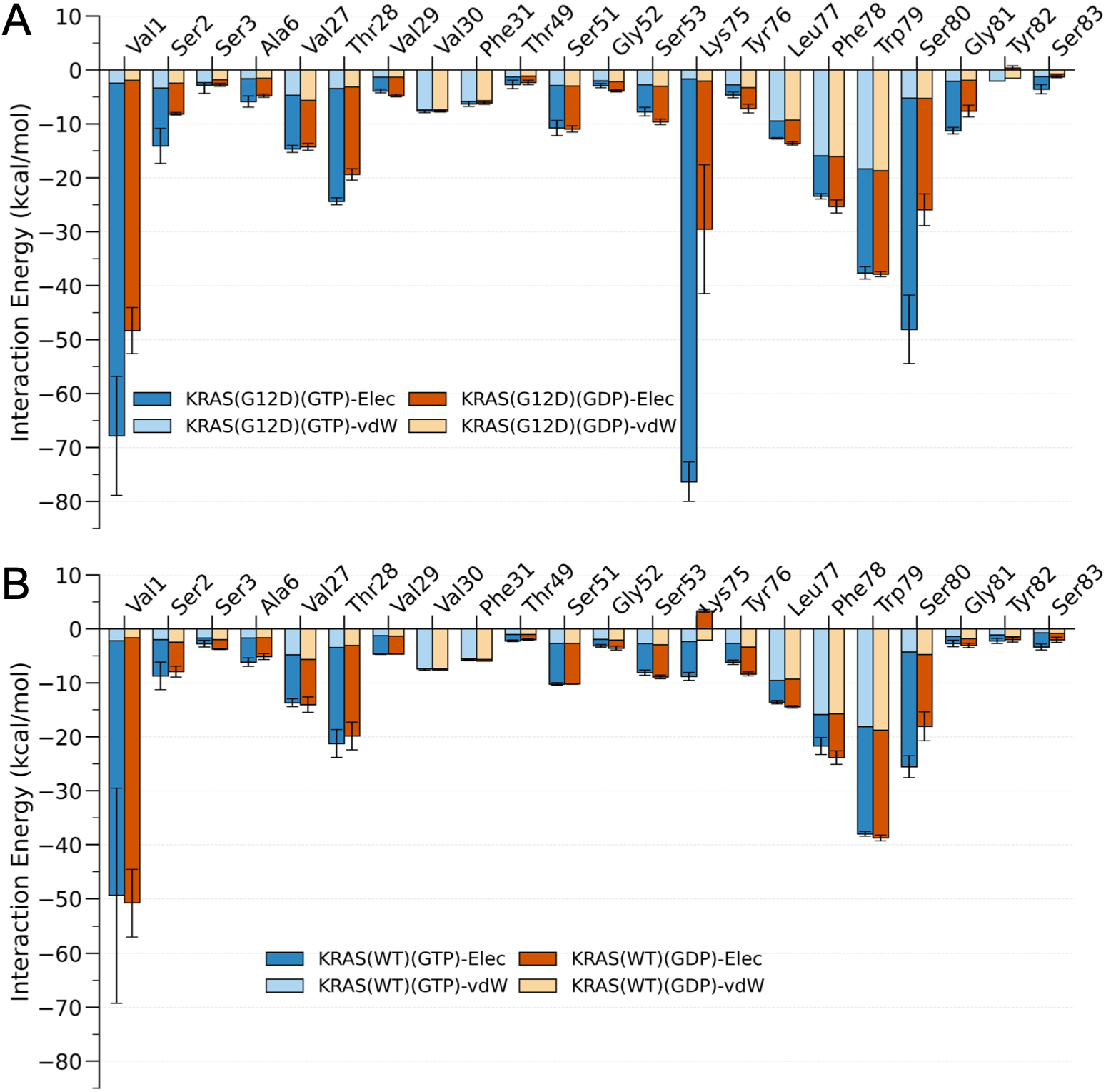
Interaction energies of 12D4 residues at the protein-protein interface upon binding to KRAS(G12D) (A) and KRAS(WT) (B) variants in GTP-bound (blue) and GDP-bound (red) states. Interaction energies were computed over the final 500 ns of KRAS-12D4 simulations. The error bars indicate the standard deviation calculated from three independent MD simulations.

**Figure 6.**
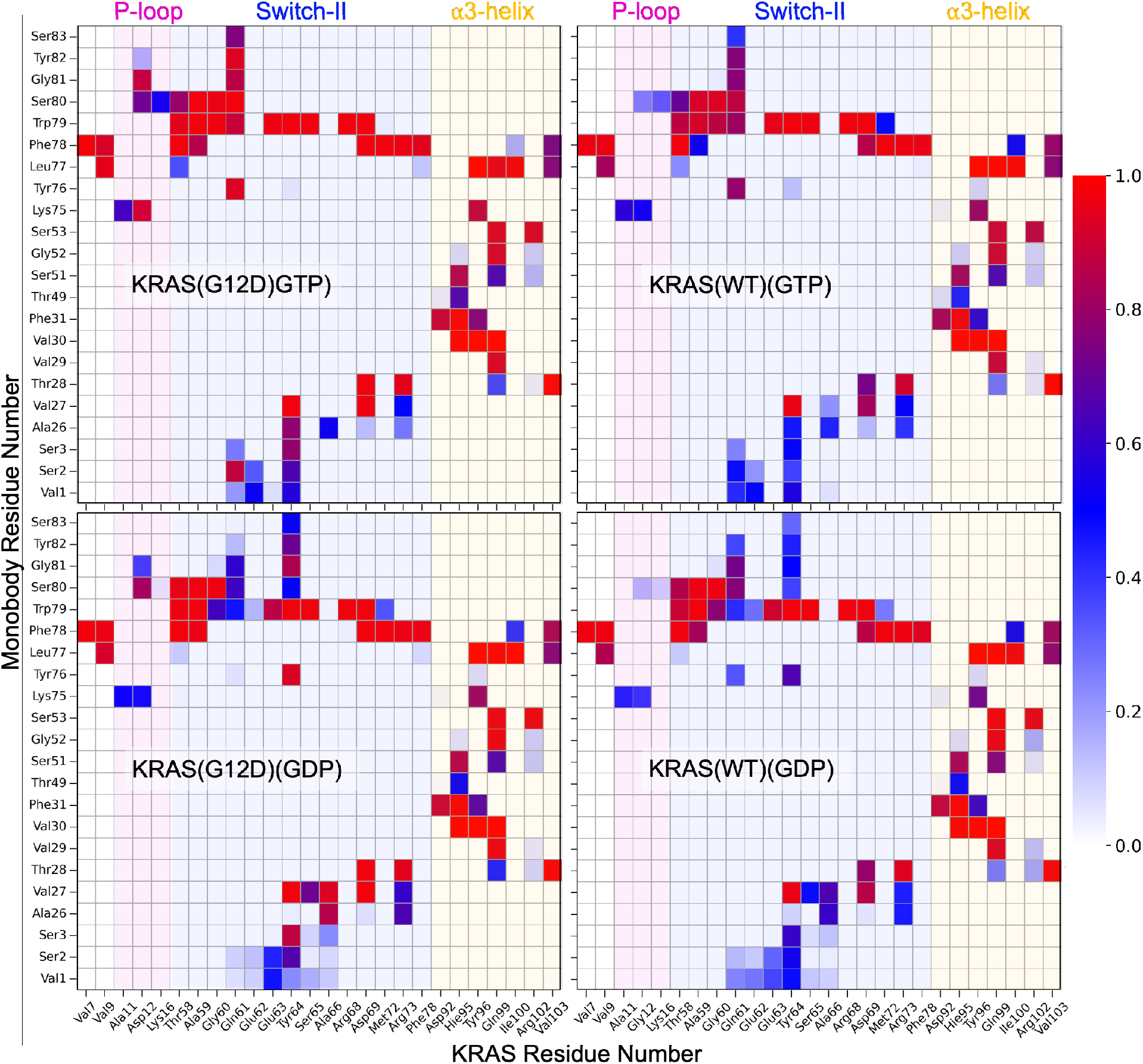
Percentage of contacts between representative residue pairs at the KRAS-12D4 interface. Contact occupancy is shown for WT and G12D variants in both GTP-and GDP-bound states. Major KRAS structural regions are colored. Contact maps for all residues at the interface are provided in Figure S4.

Comparison of different nucleotide states within each KRAS variant shows that the GTP-bound (active) state generally produces stronger interactions than the GDP-bound (inactive) state. In the G12D system, the GTP-bound state exhibits significantly enhanced electrostatic stabilization at key hotspots relative to the GDP-bound state. For example, V1 (-65.41 vs -46.46 kcal/mol), K75 (-74.71 vs -27.51 kcal/mol), and S80 (-42.86 vs -20.67 kcal/mol) show substantially stronger interactions in the GTP-bound state (**Figure 5**). These interactions are further supported by higher contact occupancies, which decrease upon transition to the GDP-bound state (**Figure 6**).

Comparison of different mutants at the same nucleotide state identifies residue 12 of KRAS as the primary determinant of binding energetics. In the GTP-bound state, G12D forms the strongest electrostatic interaction with 12D4 residue K75, followed by G12C, G12V, and WT (**Figure 5 and S5**). In contrast, the G12R mutant introduces electrostatic repulsion between KRAS R12 and 12D4 K75. In the GDP-bound state, only G12D maintains a strong interaction with K75, whereas this interaction is significantly reduced in other mutants and WT. These results identify K75 of 12D4 as a key residue for mutation-specific recognition.

Beyond these mutation-dependent hotspots, S51 in the βD strand of 12D4 contributes to maintaining stable interactions in all systems. It exhibits consistent and favorable interaction energies across all KRAS variants and nucleotide states (**Figures 5 and S5**), primarily driven by electrostatic contributions. This result suggests that S51, which replaces Pro51 in the 12D5 monobody, enhances conformational flexibility and enables sustained interactions with the KRAS α3-helix.

### Thermodynamic Integration Identifies Key Residue Determinants of Monobody Binding

To quantify the contribution of individual monobody residues to KRAS binding, we performed TI calculations for selected 12D4 mutants in complex with WT KRAS and oncogenic variants. Residues V1, V27, T28, S51, L77, F78, W79, and S80 were selected for mutagenesis because interaction energy decomposition identified them as major contributors to the binding interface. Each residue was individually mutated to Ala, except for the N-terminal residue V1. In addition, we also studied K75A mutant in the KRAS(G12D) complex because of its exceptionally favorable interaction energy.

The TI calculations identified L77, F78, and W79 within the FG loop as the primary energetic determinants of KRAS recognition (**Table 1 and Figures S7-S11**). Among these residues, F78 exhibited the largest energetic penalty upon mutation, with ΔΔG values ranging from 5.5-8.3 kcal/mol across all KRAS variants and nucleotide states. This result establishes F78 as the dominant hotspot for monobody binding. Structural analysis showed that F78 forms hydrophobic contacts with KRAS residues V7, M72, and V103 in the KRAS(G12D)(GTP) complex. Mutation of F78A disrupted the interactions with V7 and V103, which leaves only the contact with M72 (**Figure S7B**). Mutations of L77 and W79 produced smaller but still substantial energetic penalties (ΔΔG = 2-5 kcal/mol). Both residues cooperate with F78 to form a conserved hydrophobic cluster that enhances binding at the protein interface. In contrast, the S51A mutation produced minimal changes in binding energy, suggesting a limited direct contribution to interface stabilization. Note that K75 displayed a unique role in KRAS(G12D) recognition. Mutation of K75 resulted in a large increase in binding free energy in the KRAS(G12D)(GTP) complex (ΔΔG = 8.9 kcal/mol) although the effect became smaller in the GDP-bound state (ΔΔG = 3.9 kcal/mol) (**Table 1**). This strong nucleotide-state dependence suggests that the active G12D conformation promotes a favorable electrostatic interaction involving K75 that is not observed in other KRAS variants.

**Table 1.** Relative binding free energies (ΔΔG, kcal/mol) for 12D4 mutants bound to KRAS. Values represent the mean ΔΔG calculated from three independent TI calculations. Errors are standard deviation from three simulations.

|  | KRAS(G12D) |  | KRAS(G12C) |  | KRAS(G12V) |  | KRAS(G12R) |  | KRAS(WT) |  |
| --- | --- | --- | --- | --- | --- | --- | --- | --- | --- | --- |
|  | GTP | GDP | GTP | GDP | GTP | GDP | GTP | GDP | GTP | GDP |
| **12D4(V27A)** | -1.20<br>± 0.03 | 0.80 ±<br>0.21 | -0.95<br>± 0.43 | -1.32<br>± 0.51 | 1.96 ±<br>0.07 | 2.83 ±<br>0.48 | 2.10 ±<br>0.05 | 1.98 ±<br>0.22 | -1.11<br>± 0.34 | 0.70 ±<br>0.08 |
| **12D4(T28A)** | 4.38 ±<br>0.20 | 4.18 ±<br>0.20 | 1.49 ±<br>0.56 | 3.90 ±<br>0.10 | 0.48 ±<br>0.22 | 5.28 ±<br>0.30 | 3.40 ±<br>0.36 | 1.19 ±<br>0.22 | 2.27 ±<br>0.26 | 3.68 ±<br>0.25 |
| **12D4(S51A)** | 0.28 ±<br>0.02 | 0.45 ±<br>0.16 | 0.68 ±<br>0.16 | -0.20<br>± 0.36 | 0.33 ±<br>0.10 | 0.34 ±<br>0.10 | 0.13 ±<br>0.09 | 0.53 ±<br>0.10 | 0.17 ±<br>0.06 | 0.14 ±<br>0.02 |
| **12D4(K75A)** | 8.90 ±<br>1.26 | 3.90 ±<br>0.37 | NA | NA | NA | NA | NA | NA | NA | NA |
| **12D4(L77A)** | 4.24 ±<br>0.44 | 5.04 ±<br>0.40 | 0.81 ±<br>0.35 | 3.55 ±<br>0.35 | 5.23 ±<br>0.32 | 0.32 ±<br>0.33 | 4.25 ±<br>0.19 | 5.36 ±<br>0.25 | 3.17 ±<br>0.33 | 4.22 ±<br>0.52 |
| **12D4(F78A)** | 7.52 ±<br>0.56 | 7.42 ±<br>0.66 | 6.26 ±<br>0.58 | 6.47 ±<br>0.38 | 5.76 ±<br>0.49 | 5.97 ±<br>0.94 | 7.07 ±<br>0.15 | 5.08 ±<br>0.20 | 6.88 ±<br>0.65 | 8.30 ±<br>0.27 |
| **12D4(W79A)** | 4.54 ±<br>0.11 | 1.95 ±<br>0.65 | 4.36 ±<br>0.22 | 4.91 ±<br>0.62 | 7.21 ±<br>0.76 | 5.04 ±<br>0.73 | 4.03 ±<br>0.23 | 2.58 ±<br>0.41 | 4.14 ±<br>0.51 | 2.60 ±<br>0.46 |
| **12D4(S80A)** | 1.33 ±<br>0.07 | 0.87 ±<br>0.27 | 0.69 ±<br>0.06 | 0.15 ±<br>0.32 | 1.02 ±<br>0.05 | 0.43 ±<br>0.18 | 1.64 ±<br>0.32 | 1.59 ±<br>0.05 | 0.78 ±<br>0.46 | 1.33 ±<br>0.36 |

### Rational Design of 12D4 to Enhance KRAS variant Binding

Although monobody 12D4 exhibits high affinity toward KRAS(G12D), its binding to other oncogenic variants, including G12C, G12R, and G12V, is weaker. Our energy analysis identified 12D4(K75) as a key determinant of KRAS(G12D) recognition through its favorable interaction with G12D. Substitution of G12D by Cys, Arg, or Val disrupts this interaction and reduces binding affinity. To improve recognition of these variants, we computationally screened alternative amino acids at 12D4 position 75 and evaluated their effects on binding energy by TI. Several mutations produced favorable energetic effects relative to the parental K75 residue (**Figure 7A**). K75Q improved binding to both KRAS(G12C) and KRAS(G12R), whereas K75Y and K75M enhanced binding to KRAS(G12V). All selected mutants exhibited negative ΔΔG values in both GTP-and GDP-bound states, which indicates stronger binding than the original 12D4 monobody. Among them, K75Q produced the largest stabilization for KRAS(G12R), with ΔΔG values of -6.12 kcal/mol and -5.05 kcal/mol in the GTP-and GDP-bound states, respectively.

**Figure 7.**
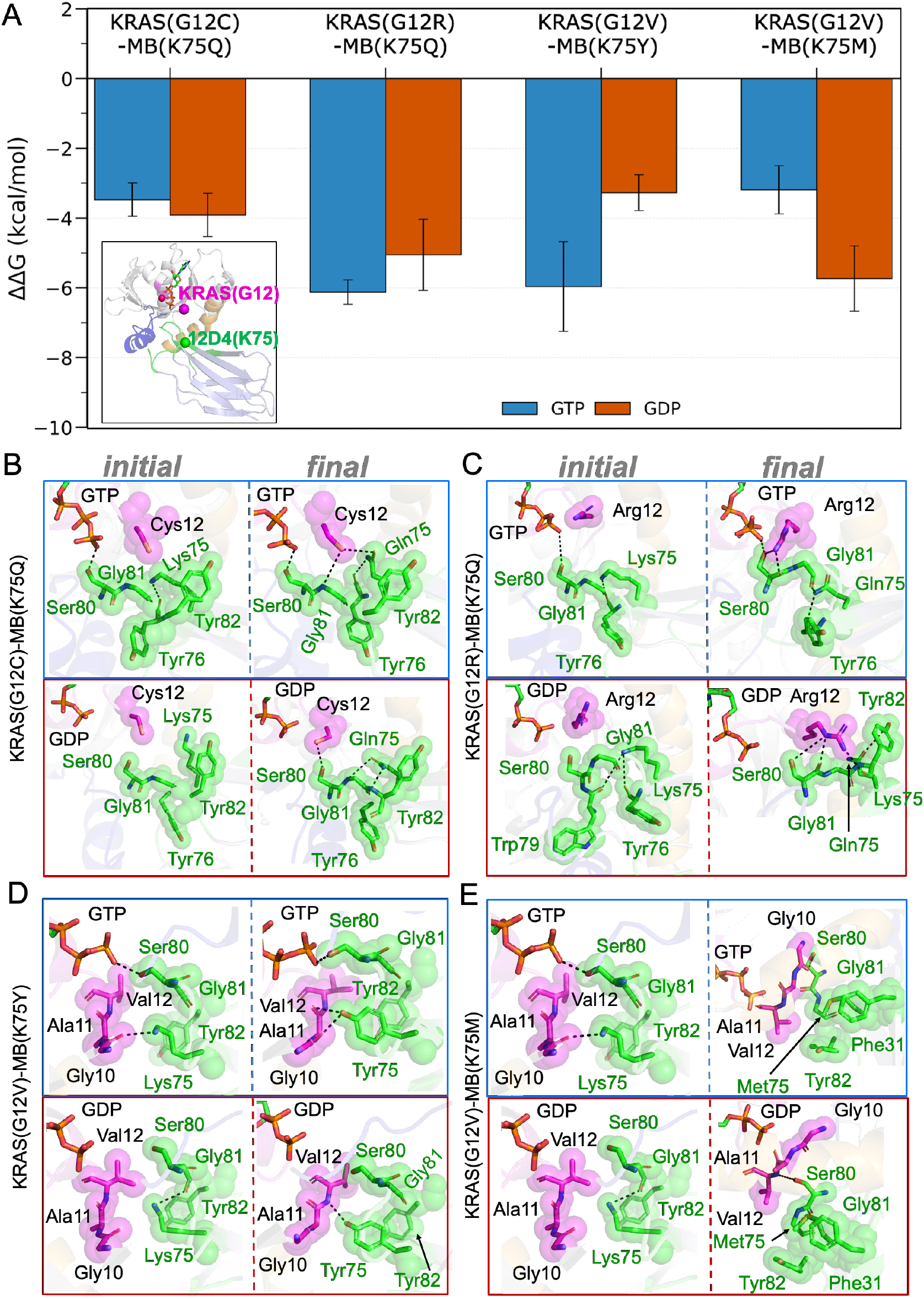
Computational design of monobody residue K75 to enhance binding toward KRAS variants. (A) Relative binding free-energy changes (ΔΔG) calculated by TI for K75 mutations in 12D4 bound to KRAS(G12C), KRAS(G12R), and KRAS(G12V) in both GTP-and GDP-bound states. Negative ΔΔG values indicate improved binding affinity relative to the parental 12D4 monobody containing K75. Error bars represent standard deviations obtained from 3 independent calculations. (B-E) Structural comparison of different 12D4 mutants. Initial and final representative structures from MD simulations are shown for (B) KRAS(G12C)-12D4(K75Q), (C) KRAS(G12R)-12D4(K75Q), (D) KRAS(G12V)-12D4(K75Y), and (E) KRAS(G12V)-12D4(K75M). KRAS residue 12 and the engineered monobody residue 75 are shown as magenta and green spheres, respectively. These mutations reorganize local interactions at the KRAS-monobody interface and promote additional favorable contacts relative to the parental K75 residue.

Structural analysis provides a mechanistic explanation for these improvements (**Figures 7B-E**). In the parental complex, K75 does not form favorable interactions with the substituted residue at KRAS position 12 and remains partially exposed to the solvent. Replacement of K75 with Gln, Tyr, or Met reorganizes the local interface and creates additional contacts with neighboring KRAS residues. For example, in the KRAS(G12C)-12D4(K75Q) complex, K75Q forms direct contacts with KRAS G12C that are absent in the parental complex (**Figure 7B**). The K75Q mutation also facilitates additional interactions between KRAS G12C and monobody residue G81, resulting in a more extensive interfacial interaction network. Similar interface rearrangements are observed in the KRAS(G12R)-12D4(K75Q) complex, where K75Q establishes favorable interactions with KRAS G12R and stabilizes the local binding environment (**Figure 7C**). In the KRAS(G12V) systems, substitutions K75Y and K75M improve packing between the monobody FG loop and the KRAS P-loop region (**Figures 7D and 7E**). These mutations increase shape complementarity at the interface and strengthen local hydrophobic contacts. These results demonstrate that residue 75 in 12D4 is a tunable hotspot for engineering monobody specificity and provide a rational strategy for developing binders with enhanced affinity toward different KRAS variants.

## Discussion

Monobodies have emerged as promising alternatives to small-molecule inhibitors for targeting KRAS because they can recognize large and dynamic protein surfaces that are difficult to access with conventional ligands. Despite recent advances in the development of KRAS-selective monobodies, the molecular determinants that govern mutant recognition and binding specificity remain incompletely understood. In this study, we combined enhanced sampling, binding energy calculations, and TI to characterize the interaction between monobody 12D4 and WT KRAS as well as several clinically relevant oncogenic variants. Our results reveal that 12D4 recognition is governed by a conserved hydrophobic interaction network centered on the FG loop of the monobody, together with mutation-specific electrostatic interactions that modulate binding affinity. These interactions influence both the conformational landscape of the KRAS switch regions and the stability of the protein-protein interface. Furthermore, computational redesign identified residue K75 as an adaptable hotspot for engineering enhanced recognition of non-G12D KRAS variants. These findings provide a molecular framework for understanding KRAS-monobody recognition and offer a rational foundation for the development of improved KRAS-targeting biologics.

The conformational changes of Switch I and Switch II in our simulations are consistent with experimental and computational evidence that KRAS exists as a dynamic ensemble rather than a single static structure^60,61^. NMR relaxation, hydrogen-deuterium exchange, and single-molecule studies show that Switch I and Switch II undergo continuous fluctuations on multiple timescales and that oncogenic mutations shift the populations of pre-existing conformational states^60^. Previous MD simulations also show that substitutions at residue 12 alter the coupling between the P-loop and switch regions, which leads to mutation-dependent changes in recognition and inhibitor accessibility^62,63^. The conformational heterogeneity in our KRAS-12D4 complexes supports this model and suggests that monobody recognition occurs through a dynamic interface rather than a rigid lock-and-key mechanism. Notably, Switch I exhibits substantial mobility, but the monobody maintains stable contacts with Switch II and the adjacent α3-helix. These contacts indicate that these regions provide a conserved recognition surface across multiple KRAS variants. This observation is consistent with structural studies of KRAS-binding monobodies, which frequently target the Switch II/α3-helix region because of its accessibility and functional importance.

Our binding energy calculations reproduce the experimentally observed preference of 12D4 for KRAS(G12D)^32^. In both nucleotide states, KRAS(G12D) exhibited the most favorable binding free energy among all systems examined and bound 12D4 approximately 5-8 kcal/mol more strongly than WT KRAS. This trend agrees well with experimental measurements, which reported binding free energies of -10.99 and -10.63 kcal/mol for KRAS(G12D) in the GTP-and GDP-bound states, respectively, compared with -7.89 and -7.82 kcal/mol for WT KRAS^32^. The corresponding dissociation constants differ by nearly two orders of magnitude, ranging from 9.8-18 nM for KRAS(G12D) and 1.8-2.0 μM for WT KRAS. The enhanced affinity of KRAS(G12D) originates from the unique electrostatic environment created by the Asp12 substitution. Unlike G12C, G12V, or the positively charged G12R mutation, G12D introduces a negatively charged side chain that can establish favorable interactions with positively charged residues at the monobody interface. The interaction between KRAS G12D and 12D4 K75 contributes major electrostatic stabilization in both nucleotide states and is largely absent in the other KRAS variants. This mechanism explains both the high affinity and remarkable selectivity reported experimentally for this monobody.

To place the 12D4 binding mechanism in a broader context, we compared its recognition mode with experimentally characterized monobodies^32,64^, affimers^65^, cyclic peptides^66,67^, and small-molecule inhibitors targeting KRAS and other RAS isoforms^7,16,29,68–70^ (**Figure 8 and Table 2**). Despite their diverse molecular structures, most high-affinity RAS binders recognize one of two major surfaces: the Switch I/II interface or the Switch II/α3 pocket. The Switch II/α3 pocket is the preferred target for most KRAS binders because it provides a well-defined allosteric site adjacent to the oncogenic mutation hotspot. Covalent ligands have also been co-crystallized in the Switch II/α3 pocket of KRAS(G12C). Here, the ligands form a covalent bond with the mutant G12C residue, thereby achieving high selectivity^71–73^. In contrast, binders developed against HRAS and NRAS more frequently engage the Switch I/II interface, where they disrupt interactions with SOS1, GAPs, and downstream effectors^68^.

**Figure 8.**
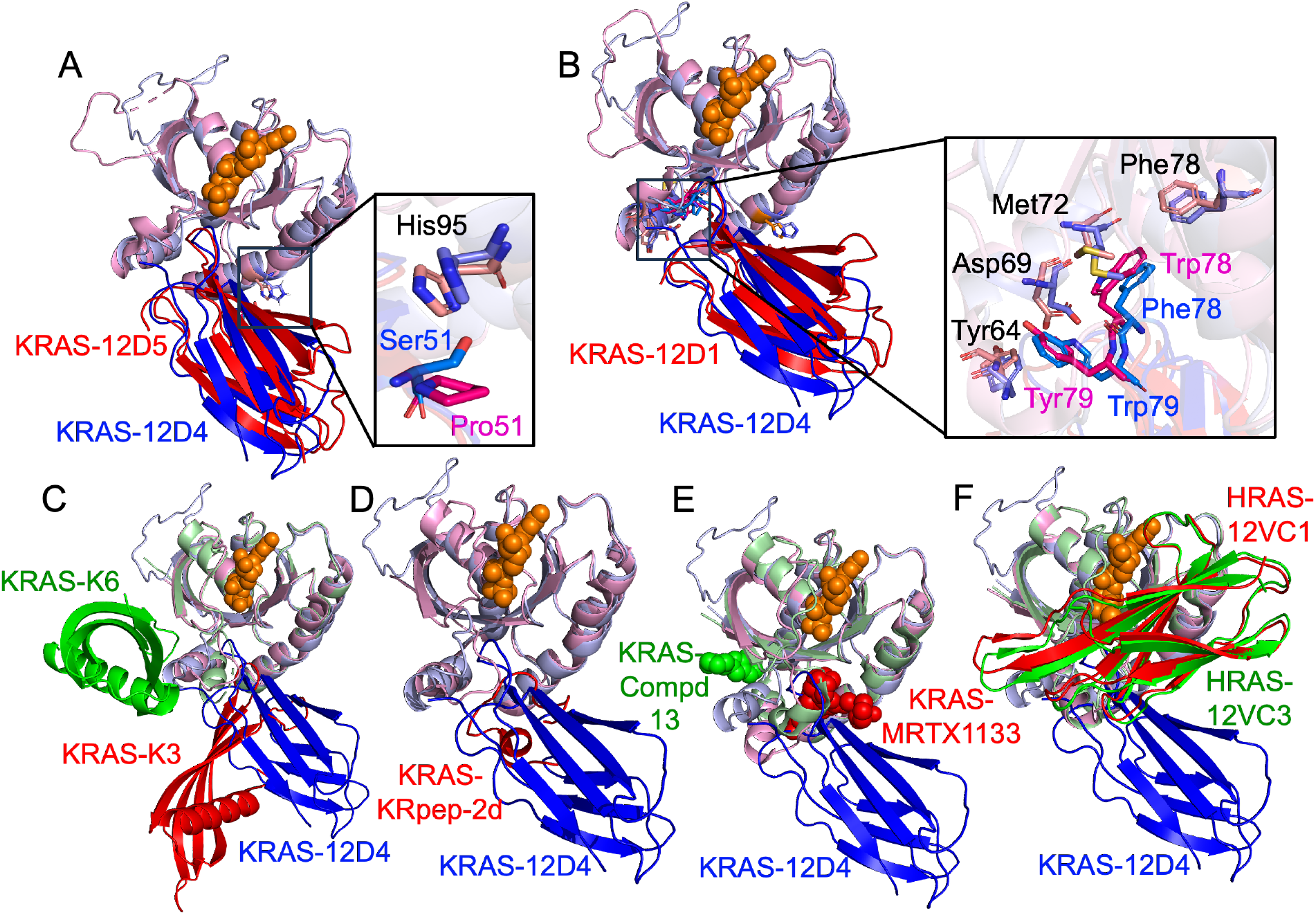
Structural comparison of the KRAS-12D4 complex with representative RAS binders. Structures were superimposed using the RAS protein to compare the binding modes of monobodies, affimers, cyclic peptides, and small-molecule inhibitors. The alignment illustrates whether each binder occupies the Switch I/II interface or the Switch II/α3-helix pocket. The KRAS-12D4 complex is shown in blue in all panels, and the bound nucleotide is represented as orange spheres. (A) Superposition with the KRAS(G12D)-12D5 complex (red; PDB ID: 8F0M). (B) Superposition with the KRAS(G12D)-12D1 complex (red; PDB ID: 8EZG). (C) Superposition with the KRAS-K3 (red; PDB ID: 6YXW) and KRASA-K6 (green; PDB ID: 6YR8) complexes.(D) Superposition with the KRAS-KRpep-2d complex (red; PDB ID: 5XCO). (E) Superposition with KRAS bound to MRTX1133 (red; PDB ID: 9NF2) and compound 13 (green; PDB ID: 6ZLI). (F) Superposition with the HRAS-12VC1 (red; PDB ID: 7L0G) and HRAS-12VC3 (green; PDB ID: 7L0F) complexes.

**Table 2.** List of experimentally determined RAS-binder complex structures deposited in the Protein Data Bank (PDB). The table summarizes the RAS isoform, mutation, corresponding protein-and small-molecule binders, binding affinity (equilibrium dissociation constants), and RAS residues that directly interact with each binder. Residues located in the P-loop, Switch I, Switch II, and α3-helix are highlighted in magenta, red, blue, and orange, respectively.

| PDB ID | Isoform | Mutation | Nucleotide State | Binder | Binder Type | Binding Affinity | KRAS Binding Site | References |
| --- | --- | --- | --- | --- | --- | --- | --- | --- |
| 8F0M | KRAS | G12D | GDP / GTPγS | 12D5 | Monobody | 108 nM / 40nM | D12, T58, A59, Q61, D69, K88, D92, H95, Y96, Q99 | PMID: 37399416 <sup>32</sup> |
| 8EZG | KRAS | G12D | GDP | 12D1 | Monobody | 720 nM | D12, A59, Q61, D69, K88, D92, H95, Q99 | PMID: 37399416 <sup>32</sup> |
| 6YR8 | KRAS | WT | GDP | K6 | Affimer | 1.36 nM | V7, I36, D38, S39, L56, Y71 | PMID: 34193876 <sup>65</sup> |
| 6YXW | KRAS | WT | GDP | K3 | Affimer | 59.4 nM | V9, E62, R68, D69, R73, M72, H95, Y96, Q99, R102, V103 | PMID: 34193876 <sup>65</sup> |
| 7L0G | HRAS | G12C | GTPγS | 12VC1 | Monobody | 24.7 nM | C12, D33, P34, Q61, K117 | PMID: 33976200 <sup>64</sup> |
| 7L0F | HRAS | WT | GTPγS | 12VC3 | Monobody | NA | G12, D33, P34, E63, N85, K117 | PMID: 33976200 <sup>33</sup> |
| 6WGN | KRAS | G12D | GppNHp | KD2 | Cyclic Peptide | NA | D12, G60, D69, D92, Y96, Q99 | PMID: 33145412 <sup>67</sup> |
| 5XCO | KRAS | G12D | GDP | KRpep-2d | Cyclic Peptide | 8.9 nM | V9, D12, T58, Q61, R68, D69, M72, R73, Y96, Q99, R102, V103 | PMID: 28740607 <sup>66</sup> |
| 7RPZ | KRAS | G12D | GDP | MRTX1133 | Small Molecule | 2 nM | V9, D12, G60, E62, Y64, R68, D69, H95, Y96, Q99 | PMID: 34889605 <sup>69</sup> |
| 7RT1 | KRAS | G12D | GDP | Compound 15 | Small Molecule | 0.8 mM | G10, D12, G60, E62, Y64, D69, H95, Y96 | PMID: 34889605 <sup>69</sup> |
| 7RT2 | KRAS | G12D | GDP | Compound 25 | Small Molecule | 2 mM | D12, G60, D62, Y64, R68, D69, H95, Y96 | PMID: 34889605 <sup>69</sup> |
| 7RT3 | KRAS | G12D | GDP | Compound 24 | Small Molecule | 2 mM | D12, G60, Y64, D69, M72, H95, Y96 | PMID: 34889605 <sup>69</sup> |
| 7RT4 | KRAS | G12D | GDP | Compound 5B | Small Molecule | 3.5 μM | D12, G60, Y64, D69, M72, H95, Y96 | PMID: 34889605 <sup>69</sup> |
| 7RT5 | KRAS | G12D | GDP | Compound 36 | Small Molecule | 2 mM | G10, D12, E62, Y64, R68, H95, Y96 | PMID: 34889605 <sup>69</sup> |
| 9NF2 | KRAS | G12D | GMPPNP | MRTX1133 | Small Molecule | NA | D12, G60, Y64, R68, M72, H95, Y96, R102 | NA |
| 9O0R | KRAS | WT | GDP | MRTX849 | Small Molecule | NA | G10, G12, K16, E62, Y64, M72 H95, Y96 | PMID: 40473215 <sup>70</sup> |
| 6ZLI | KRAS | G12D | GCP | Compound 13 | Small Molecule | 1.9 mM | K5, D54, L56 | PMID: 32779487 <sup>68</sup> |
| 6ZIR | NRAS | C118S | GDP | Compound 18 | Small Molecule | NA | E37, D54, L56, T74 | PMID: 32779487 <sup>68</sup> |
| 6ZIZ | NRAS | Q61R | GTP | Compound 18 | Small Molecule | 33 μM | K5, E37, D54, L56 | PMID: 32779487 <sup>68</sup> |
| 6ZJ0 | HRAS | G12D | GCP | Compound 18 | Small Molecule | 6.7 μM | K5, E37, D54, L56, T74 | PMID: 32779487 <sup>68</sup> |
| 6ZL3 | HRAS | WT | GDP | Compound 18 | Small Molecule | 54 μM | K5, E37, D54, L56, T74 | PMID: 32779487 <sup>68</sup> |
| 6ZL5 | KRAS | G12D/C118S | GDP | BI-2852 | Small Molecule | 2 μM | K5, E38, S39, D54, L56, T74 | PMID: 32779487 <sup>68</sup> |
| 6OIM | KRAS | G12C | GDP | AMG 510 | Small Molecule | NA | C12, K30, Q75, E76, E77, H95, Y96 Q99 | PMID: 31666701 <sup>74</sup> |
| 4LUC | KRAS | G12C | GDP | Compound 6 | Small Molecule | NA | V10, C12, G61, Y97 | PMID: 24256730 <sup>75</sup> |
| 9E9H | KRAS | G12C | GDP | Compound 5 | Small Molecule | NA | C12, K16, R68, H95, Y96, Q99 | PMID: 39930787 <sup>72</sup> |

The binding mode of 12D4 closely resembles that of the KRAS(G12D)-selective monobodies 12D1 and 12D5^32^. In all three systems, the monobody FG loop inserts into the Switch II/α3 pocket and establishes extensive contacts with residues from Switch II and the α3-helix. However, subtle sequence differences produce measurable changes in affinity. Our simulations suggest that S51 in 12D4 contributes favorable interactions with H95 on the KRAS α3-helix, whereas the corresponding P51 residue in 12D1 and 12D5 lacks this interaction (**Figure 8A and 8B**). Additional differences occur within the FG loop. The F78-W79 pair in 12D4 adopts a geometry that complements the Switch II/α3 pocket, whereas the corresponding W78-Y79 residues in 12D1 generate less favorable packing interactions (**Figure 8B**). These findings are consistent with experimental measurements showing that 12D4 binds KRAS(G12D) more strongly than either 12D1 or 12D5. Furthermore, the direct interaction between 12D4 residue K75 and KRAS residue G12D (**Figure 7**) provides an additional source of mutant-specific stabilization that is absent from the available crystal structures of 12D1 and 12D5. These residue-level interactions also provide practical guidance for future RAS-targeting binder design. The K75-G12D interaction highlights the value of positioning charged residues near the mutation site to enhance mutation-specific recognition, while optimization of the FG-loop residues may improve complementarity within the Switch II/α3 pocket. This strategy could be extended to other RAS variants by identifying adaptable monobody residues that form favorable interactions with the distinct chemical environments created by oncogenic mutations.

A similar recognition strategy is found in several non-monobody KRAS binders. The affimer K3^65^, the cyclic peptide KRpep-2d^66^, and the MRTX1133 series^69^ (e.g., compound 15, 25, 24, 5B, and 36) all target the Switch II/α3 pocket and exploit interactions near the G12D mutation site (**Figure 8C, 8D, and 8E**). Among these binders, MRTX1133 achieves particularly high affinity through extensive contacts with residues surrounding G12D and efficient occupation of the hydrophobic Switch II subpocket^69^. In contrast, affimer K6^65^, BI-2852 derivatives^68^ (e.g., compound 13 and 18) (**Figure 8E**), and the monobodies 12VC1 and 12VC3^33^ (**Figure 8F**) bind primarily at the Switch I/II interface. These molecules inhibit RAS signaling by blocking interactions with downstream effectors rather than directly exploiting the Switch II/α3 allosteric network. These comparisons suggest that the Switch II/α3 pocket represents a privileged site for achieving high-affinity and mutation-selective recognition of KRAS, while optimization of local contacts near residue 12 provides an effective strategy for enhancing mutant specificity.

## Data, Materials, and Software Availability

The simulation input files and resulting trajectories are freely available at https://doi.org/10.5281/zenodo.21140181.

## Supporting Information

Additional structural, conformational, and residue-level interaction analyses of the KRAS-monobody systems.

## Acknowledgments

We thank the Wayne State University (WSU) High Performance Computing Center for providing the resources used for GaMD simulations and post-MD analysis. We also thank Tushan Gamage for proofreading the manuscript and providing helpful comments on the text. This work was supported by WSU startup funds, WSU Postdoctoral Fellows funds, and NIH grant R35GM160192 (to Y.M.H.).

## TOC

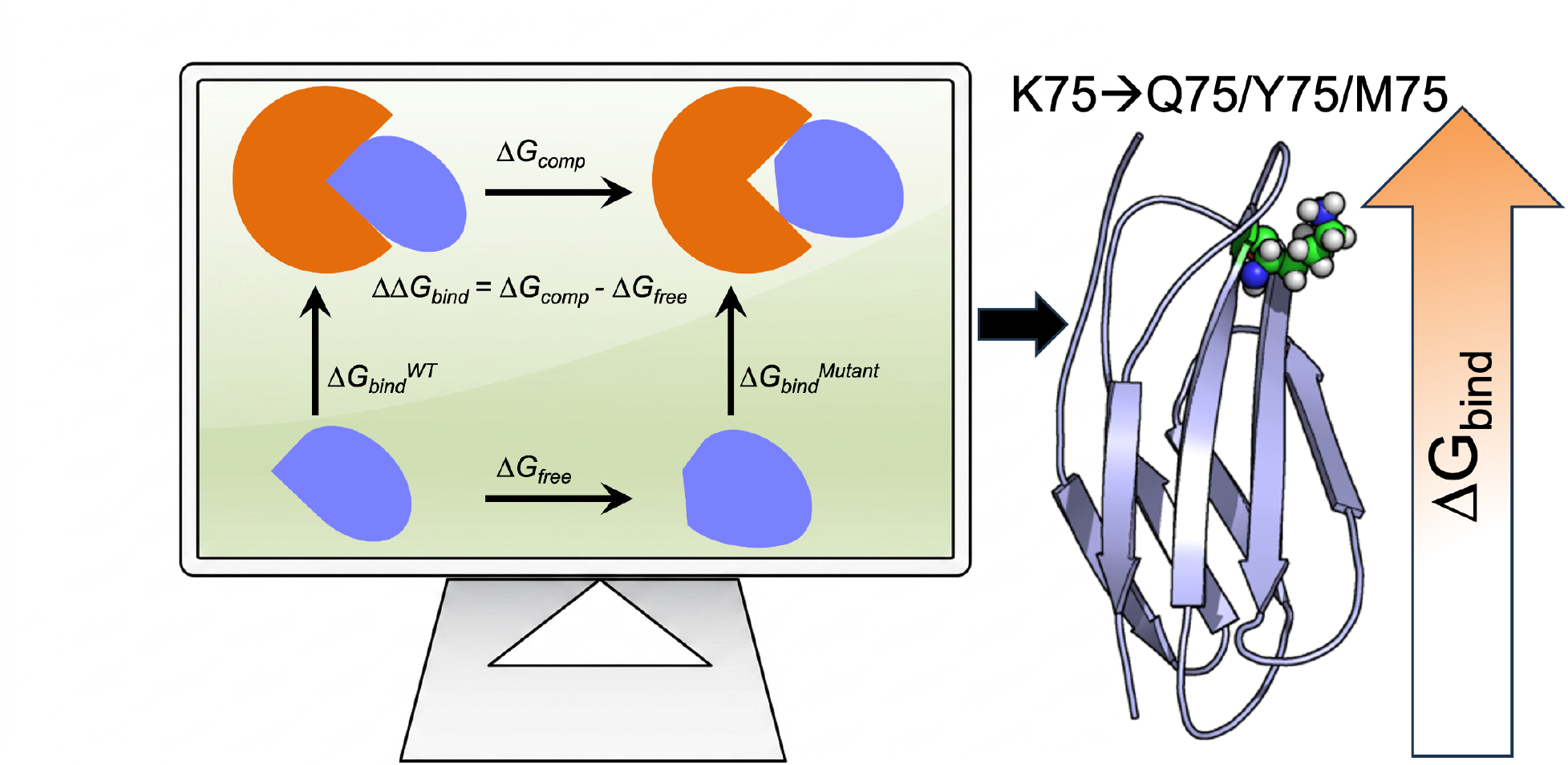

