## Supplementary Information for "Structural and Energetic Determinants of Monobody Recognition of Oncogenic KRAS Variants"

| KRAS-GTP Complex | KRAS | Monobody | P-loop | Switch I | Switch II | α3-helix |
|---|---|---|---|---|---|---|
| **G12D** | 3.79 ± 0.46 | 2.23 ± 0.26 | 0.22 ± 0.03 | 3.30 ± 0.39 | 0.54 ± 0.08 | 0.52 ± 0.07 |
| **G12C** | 4.42 ± 0.32 | 2.13 ± 0.26 | 0.24 ± 0.03 | 3.38 ± 0.38 | 1.03 ± 0.22 | 0.76 ± 0.10 |
| **G12R** | 2.70 ± 0.49 | 1.85 ± 0.22 | 0.29 ± 0.03 | 3.03 ± 0.68 | 1.04 ± 0.21 | 0.87 ± 0.10 |
| **G12V** | 3.74 ± 0.26 | 2.08 ± 0.26 | 0.32 ± 0.04 | 3.87 ± 0.27 | 0.96 ± 0.13 | 0.64 ± 0.08 |
| **WT** | 3.05 ± 0.57 | 1.96 ± 0.32 | 0.31 ± 0.05 | 2.79 ± 0.57 | 0.67 ± 0.18 | 0.71 ± 0.08 |

**Table S1:** Average root-mean-square deviation (RMSD, Å) values for KRAS(GTP)-monobody 12D4 complexes across different variants. RMSD values are reported for the full KRAS protein, the monobody, and key KRAS structural elements: P-loop, Switch I, Switch II, and α3 helix.

| KRAS-GDP Complex | KRAS | Monobody | P-loop | Switch I | Switch II | α3-helix |
|---|---|---|---|---|---|---|
| **G12D** | 3.13 ± 0.45 | 1.65 ± 0.18 | 0.26 ± 0.03 | 3.19 ± 0.26 | 1.25 ± 0.26 | 0.64 ± 0.09 |
| **G12C** | 3.02 ± 0.30 | 2.09 ± 0.28 | 0.26 ± 0.04 | 2.72 ± 0.29 | 0.95 ± 0.12 | 0.61 ± 0.08 |
| **G12R** | 4.09 ± 0.38 | 1.84 ± 0.21 | 0.28 ± 0.04 | 3.44 ± 0.31 | 1.10 ± 0.21 | 0.71 ± 0.09 |
| **G12V** | 2.31 ± 0.28 | 2.14 ± 0.27 | 0.25 ± 0.03 | 2.90 ± 0.50 | 1.29 ± 0.37 | 0.84 ± 0.10 |
| **WT** | 4.06 ± 0.44 | 1.96 ± 0.22 | 0.29 ± 0.04 | 2.86 ± 0.33 | 1.10 ± 0.14 | 0.51 ± 0.07 |

**Table S2:** Average root-mean-square deviation (RMSD, Å) values for KRAS(GDP)-monobody 12D4 complexes across different variants. RMSD values are reported for the full KRAS protein, the monobody, and key KRAS structural elements: P-loop, Switch I, Switch II, and α3 helix.

| KRAS-GTP Complex | $\Delta E_{vdW}$ | $\Delta E_{coul}$ | $\Delta W_{PB}$ | $\Delta W_{np}$ | $\Delta W_{disp}$ | $\Delta G_{bind}$ |
|---|---|---|---|---|---|---|
| **G12D** | -111.18 ± 1.84 | -52.15 ± 3.57 | 62.20 ± 2.94 | -79.78 ± 1.72 | 147.24 ± 2.53 | -33.68 ± 2.67 |
| **G12C** | -109.24 ± 3.60 | -29.63 ± 2.85 | 41.90 ± 3.18 | -76.54 ± 2.16 | 141.51 ± 3.14 | -32.02 ± 2.99 |
| **G12R** | -109.52 ± 2.71 | -23.21 ± 6.05 | 35.36 ± 5.24 | -76.39 ± 2.28 | 140.40 ± 3.23 | -33.37 ± 2.71 |
| **G12V** | -109.07 ± 2.9 | -31.54 ± 6.14 | 44.45 ± 6.46 | -76.67 ± 1.89 | 141.20 ± 3.51 | -31.64 ± 2.47 |
| **WT** | -103.40 ± 2.21 | -29.79 ± 2.79 | 42.08 ± 1.50 | -72.73 ± 1.96 | 135.76 ± 2.73 | -28.09 ± 2.80 |

**Table S3:** Energy (kcal/mol) obtained from MM/PBSA calculations. The energy is computed according to the final 500-ns simulations of KRAS-12D4 complexes in the GTP-bound active KRAS state. $\Delta E_{vdW}$ and $\Delta E_{coul}$ are the van der Waals and electrostatic interactions, respectively. $\Delta W_{PB}$ is the polar solvation free energy calculated by the Poisson-Boltzmann model. $\Delta W_{np}$ is the nonpolar contribution to create a cavity in the water. $\Delta W_{disp}$ represents the contribution of dispersion interaction between the solute and the solvent. The systems mentioned in the first column indicate the different variants of the KRAS in the GTP bound active state.

| KRAS-GDP Complex | $\Delta E_{vdw}$ | $\Delta E_{coul}$ | $\Delta W_{PB}$ | $\Delta W_{np}$ | $\Delta W_{disp}$ | $\Delta G_{bind}$ |
|---|---|---|---|---|---|---|
| **G12D** | -112.65 ± 1.27 | -31.30 ± 3.81 | 42.29 ± 3.13 | -78.93 ± 1.62 | 144.84 ± 1.79 | -35.75 ± 2.07 |
| **G12C** | -106.13 ± 4.95 | -25.62 ± 3.06 | 38.06 ± 2.77 | -74.26 ± 3.11 | 136.75 ± 5.25 | -31.20 ± 3.34 |
| **G12R** | -96.10 ± 3.36 | -25.05 ± 5.09 | 35.22 ± 2.42 | -66.52 ± 2.60 | 124.39 ± 3.64 | -28.07 ± 4.86 |
| **G12V** | -102.75 ± 0.54 | -26.37 ± 4.99 | 38.16 ± 4.41 | -71.60 ± 1.28 | 134.07 ± 0.96 | -28.50 ± 2.38 |
| **WT** | -106.56 ± 2.91 | -29.64 ± 2.80 | 40.64 ± 2.47 | -74.27 ± 1.96 | 137.29 ± 3.08 | -32.55 ± 2.20 |

**Table S4:** Energy (kcal/mol) obtained from MM/PBSA calculations. The energy is computed according to the final 500-ns simulations of KRAS-12D4 complexes in the GDP-bound inactive KRAS state. $\Delta E_{vdw}$ and $\Delta E_{coul}$ are the van der Waals and electrostatic interactions, respectively. $\Delta W_{PB}$ is the polar solvation free energy calculated by the Poisson-Boltzmann model. $\Delta W_{np}$ is the nonpolar contribution to create a cavity in the water. $\Delta W_{disp}$ represents the contribution of dispersion interaction between the solute and the solvent. The systems mentioned in the first column indicate the different variants of the KRAS in the GDP bound inactive state.

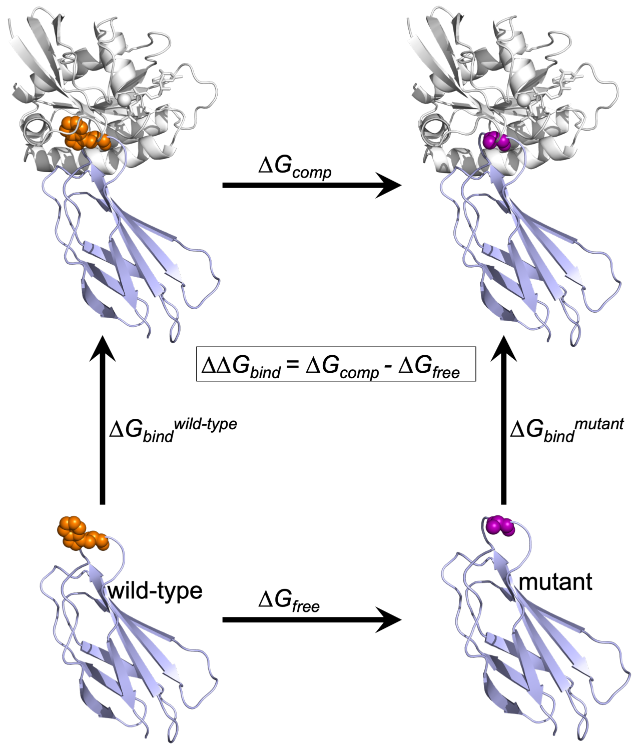


**Figure S1:** The thermodynamic cycle used in estimating the relative binding free energy of a mutation in 12D4 monobody. 12D4 and KRAS are shown in purple and white cartoons, respectively. The vertical arms of the thermodynamic cycle correspond to KRAS binding to WT and mutated monobody 12D4. The horizontal arms represent the free energy changes associated with a mutation (e.g., Trp to Ala) in the KRAS-12D4 complex (top panels) and in the monobody (bottom panels).

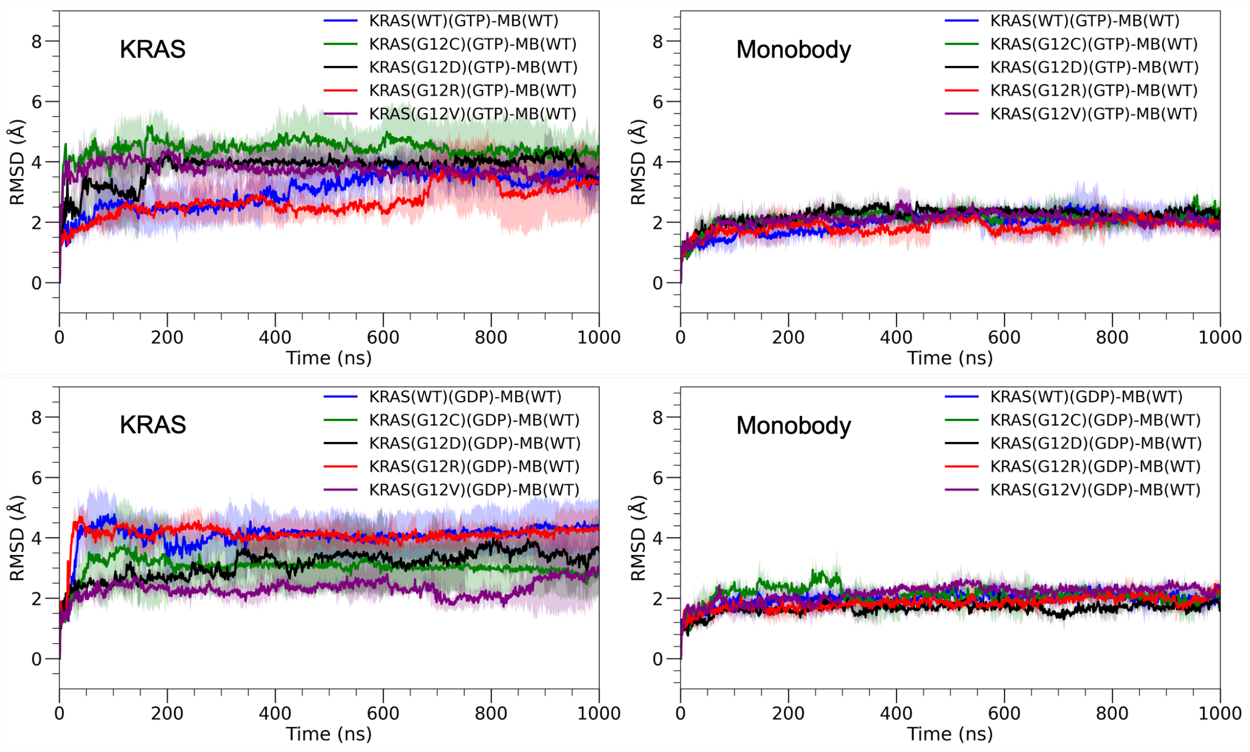


**Figure S2:** Time evolution of root-mean-square deviation (RMSD) for five KRAS systems in complex with the monobody 12D4, shown for both active (GTP-bound) and inactive (GDP-bound) states.

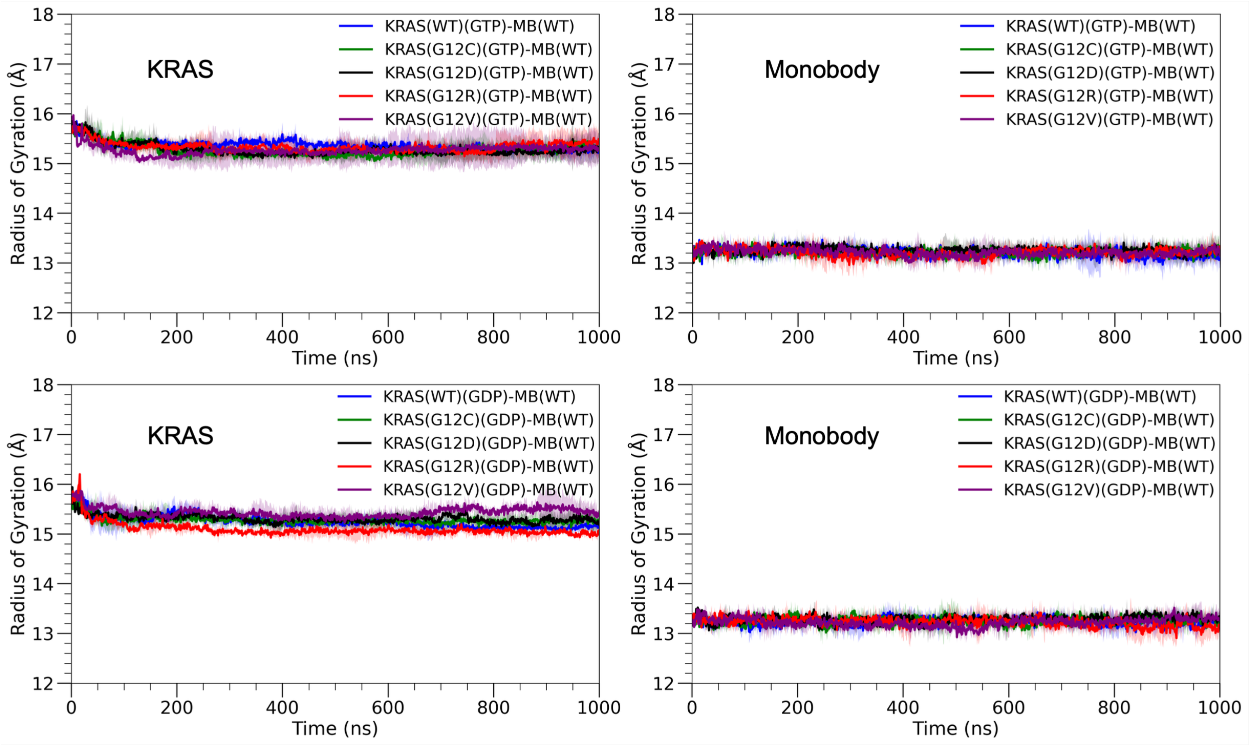


**Figure S3:** Time evolution of the radius of gyration (Rg) for five KRAS systems in complex with the monobody 12D4, shown for both active (GTP-bound) and inactive (GDP-bound) states.

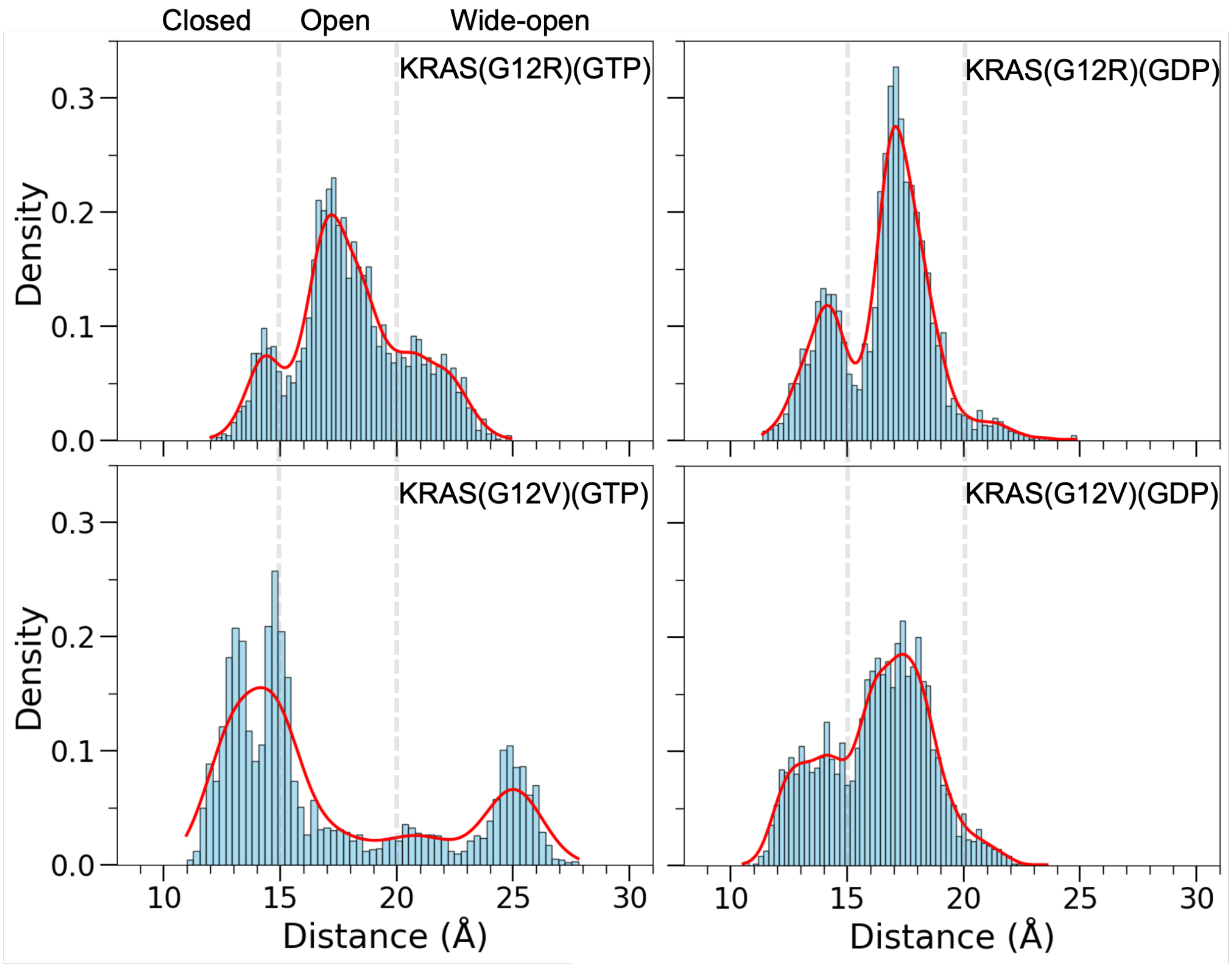


**Figure S4:** Probability density distributions show the center-of-mass distances between Switch I and Switch II for KRAS(G12R) and KRAS(G12V) in GTP- and GDP-bound states. Vertical dashed lines indicate the boundaries between closed (<15 Å), open (15-20 Å), and wide-open (>20 Å) conformational states.

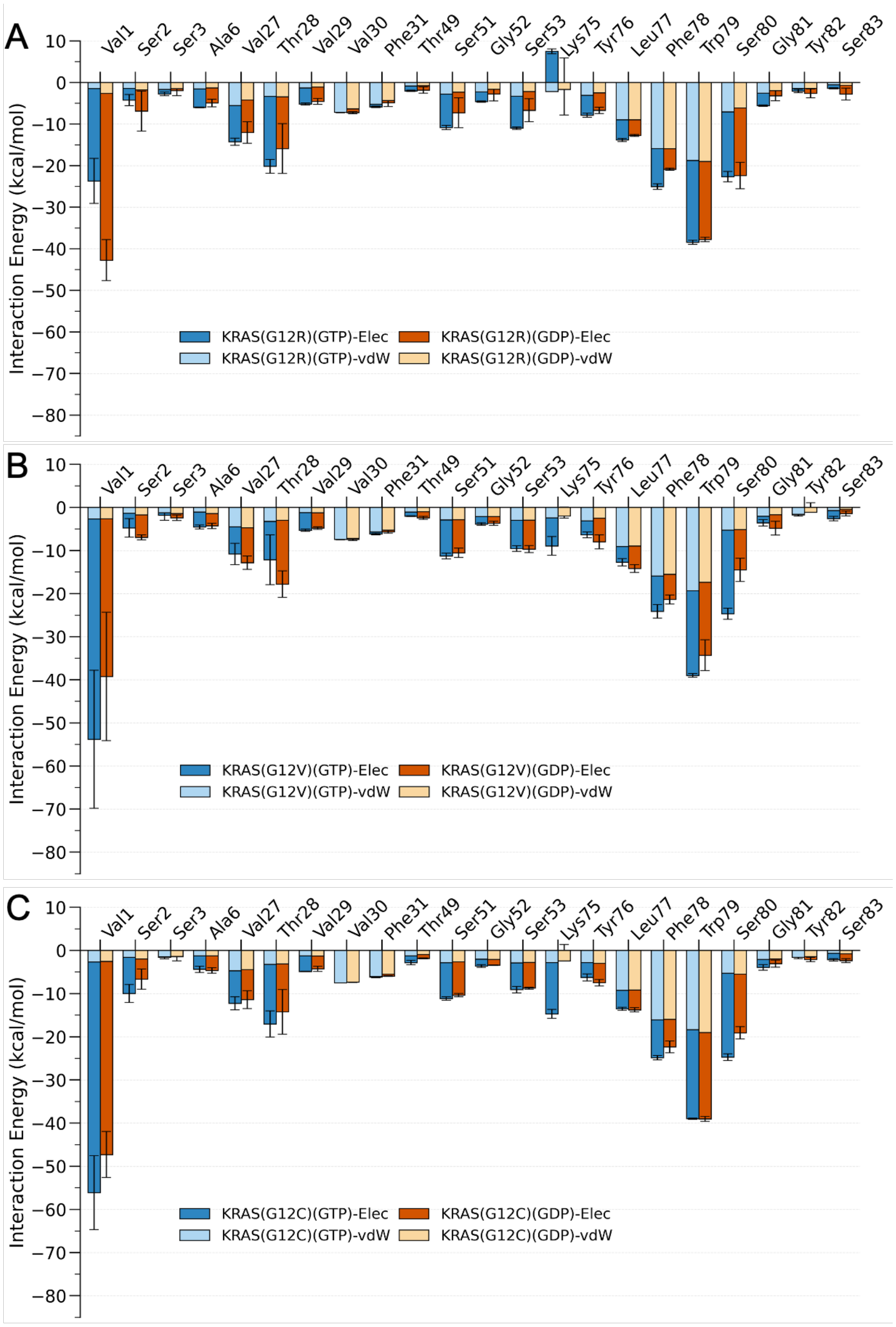


**Figure S5**: Interaction energies of 12D4 residues at the protein-protein interface upon binding to KRAS G12R (A), G12V (B), and G12C (C) variants in GTP-bound (blue) and GDP-bound (red) states. Interaction energies are computed over the final 500 ns of KRAS-12D4 simulations. The error bars indicate the standard deviation calculated from three independent MD simulations.

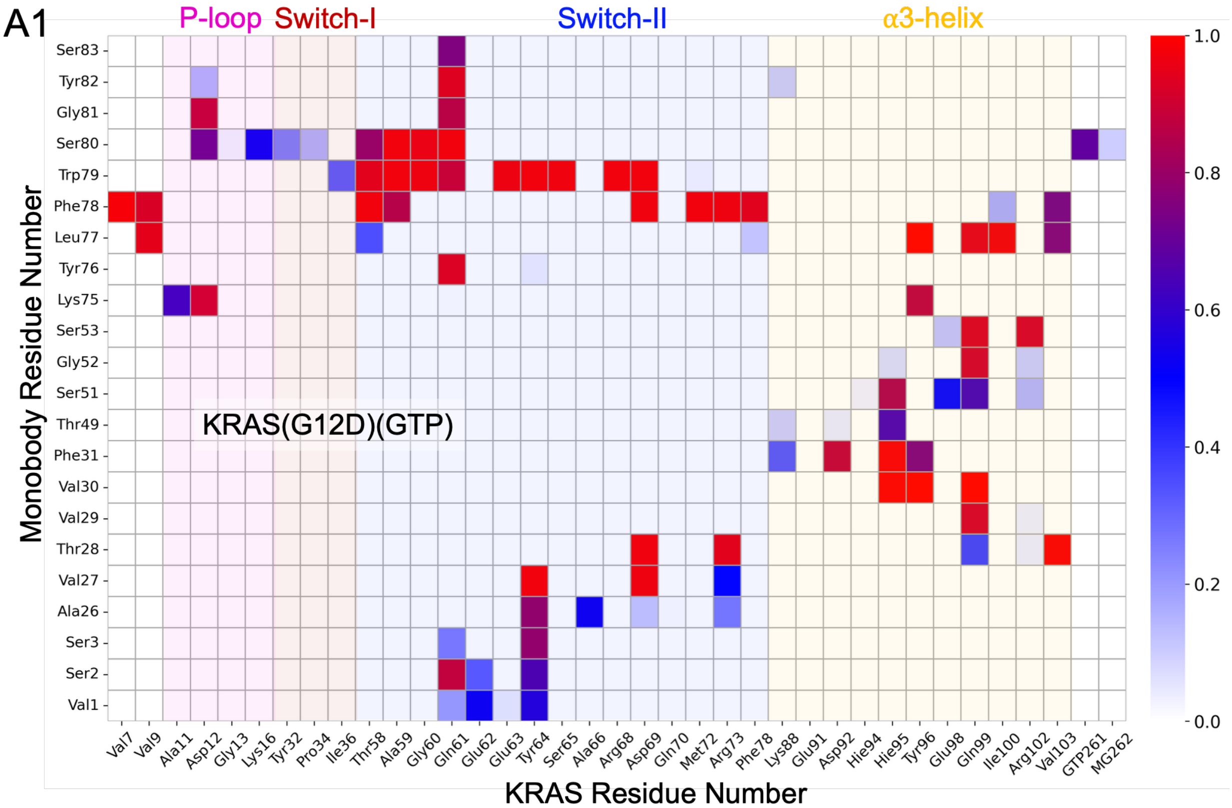
A1
P-loop
Switch-I
Switch-II
α3-helix
KRAS(G12D)(GTP)
Monobody Residue Number
Ser83
Tyr82
Gly81
Ser80
Trp79
Phe78
Leu77
Tyr76
Lys75
Ser53
Gly52
Ser51
Thr49
Phe31
Val30
Val29
Thr28
Val27
Ala26
Ser3
Ser2
Val1
Val7 Val9 Ala11 Asp12 Gly13 Lys16 Tyr32 Pro34 Ile36 Thr58 Ala59 Gly60 Gln61 Glu62 Glu63 Tyr64 Ser65 Ala66 Arg68 Asp69 Gln70 Met72 Arg73 Phe78 Lys88 Glu91 Asp92 Hie94 Hie95 Tyr96 Glu98 Gln99 Ile100 Arg102 Val103 GTP261 MG262
KRAS Residue Number
1.0
0.8
0.6
0.4
0.2
0.0

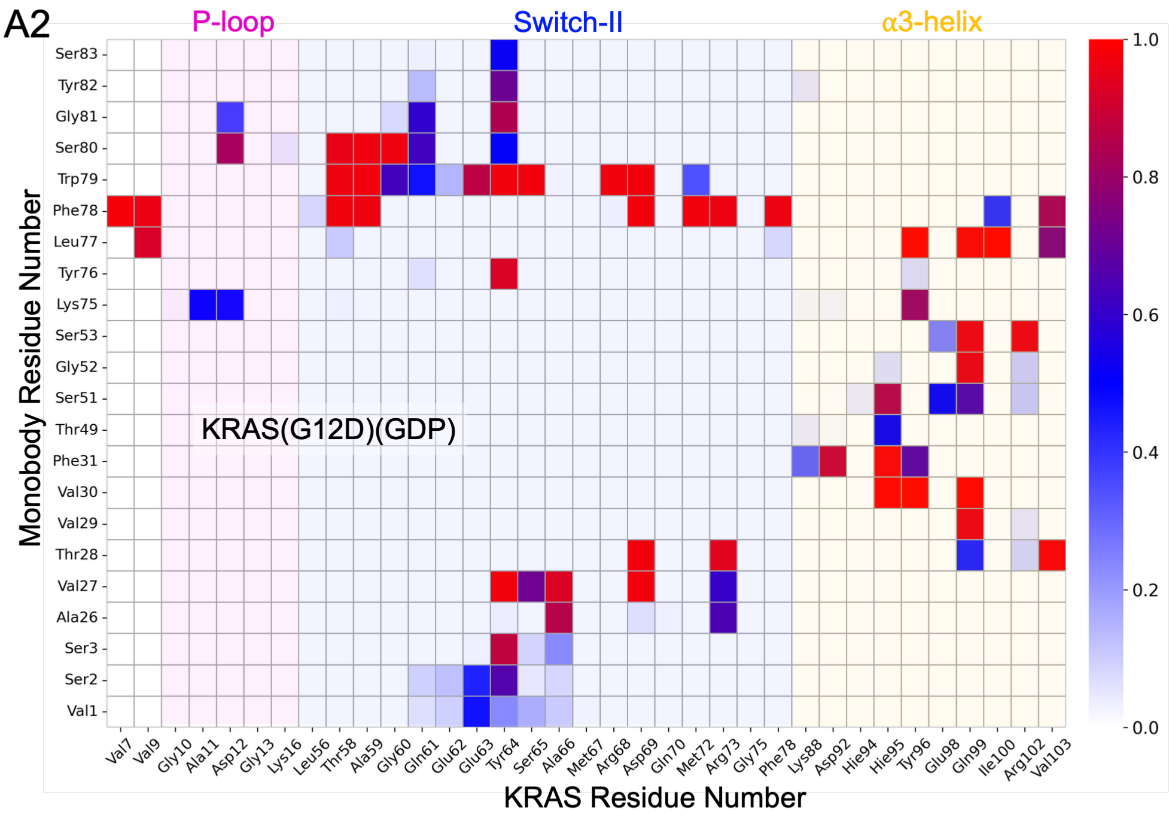
A2
P-loop
Switch-II
α3-helix
KRAS(G12D)(GDP)
Monobody Residue Number
Ser83
Tyr82
Gly81
Ser80
Trp79
Phe78
Leu77
Tyr76
Lys75
Ser53
Gly52
Ser51
Thr49
Phe31
Val30
Val29
Thr28
Val27
Ala26
Ser3
Ser2
Val1
Val7 Val9 Gly10 Ala11 Asp12 Gly13 Lys16 Leu56 Thr58 Ala59 Gly60 Gln61 Glu62 Glu63 Tyr64 Ser65 Ala66 Met67 Arg68 Asp69 Gln70 Met72 Arg73 Gly75 Phe78 Lys88 Asp92 Hie94 Hie95 Tyr96 Glu98 Gln99 Ile100 Arg102 Val103
KRAS Residue Number
1.0
0.8
0.6
0.4
0.2
0.0

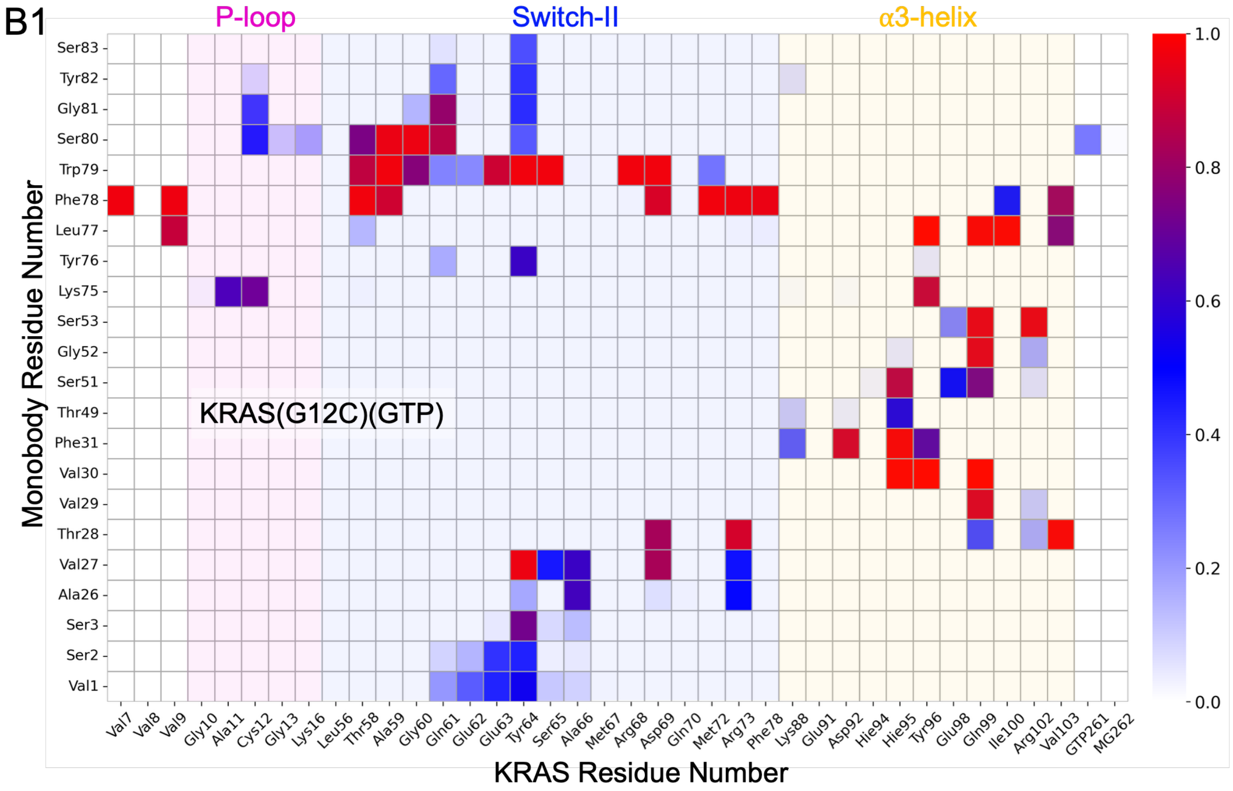
B1
P-loop
Switch-II
α3-helix
KRAS(G12C)(GTP)
Monobody Residue Number
KRAS Residue Number
Ser83
Tyr82
Gly81
Ser80
Trp79
Phe78
Leu77
Tyr76
Lys75
Ser53
Gly52
Ser51
Thr49
Phe31
Val30
Val29
Thr28
Val27
Ala26
Ser3
Ser2
Val1
Val7
Val8
Val9
Gly10
Ala11
Cys12
Gly13
Lys16
Leu56
Thr58
Ala59
Gly60
Gln61
Glu62
Glu63
Tyr64
Ser65
Ala66
Met67
Arg68
Asp69
Gln70
Met72
Arg73
Phe78
Lys88
Glu91
Asp92
Hie94
Hie95
Tyr96
Glu98
Gln99
Ile100
Arg102
Val103
GTP261
MG262
1.0
0.8
0.6
0.4
0.2
0.0

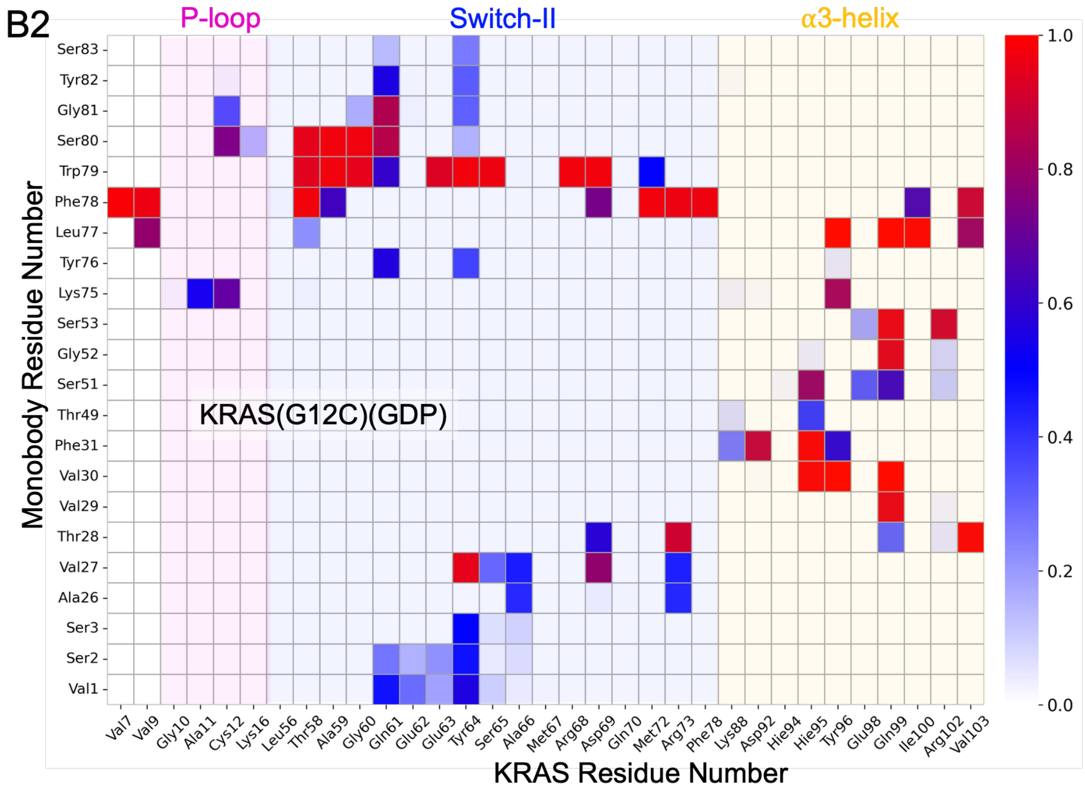
B2
P-loop
Switch-II
α3-helix
KRAS(G12C)(GDP)
Monobody Residue Number
KRAS Residue Number
Ser83
Tyr82
Gly81
Ser80
Trp79
Phe78
Leu77
Tyr76
Lys75
Ser53
Gly52
Ser51
Thr49
Phe31
Val30
Val29
Thr28
Val27
Ala26
Ser3
Ser2
Val1
Val7
Val9
Gly10
Ala11
Cys12
Lys16
Leu56
Thr58
Ala59
Gly60
Gln61
Glu62
Glu63
Tyr64
Ser65
Ala66
Met67
Arg68
Asp69
Gln70
Met72
Arg73
Phe78
Lys88
Asp92
Hie94
Hie95
Tyr96
Glu98
Gln99
Ile100
Arg102
Val103
1.0
0.8
0.6
0.4
0.2
0.0

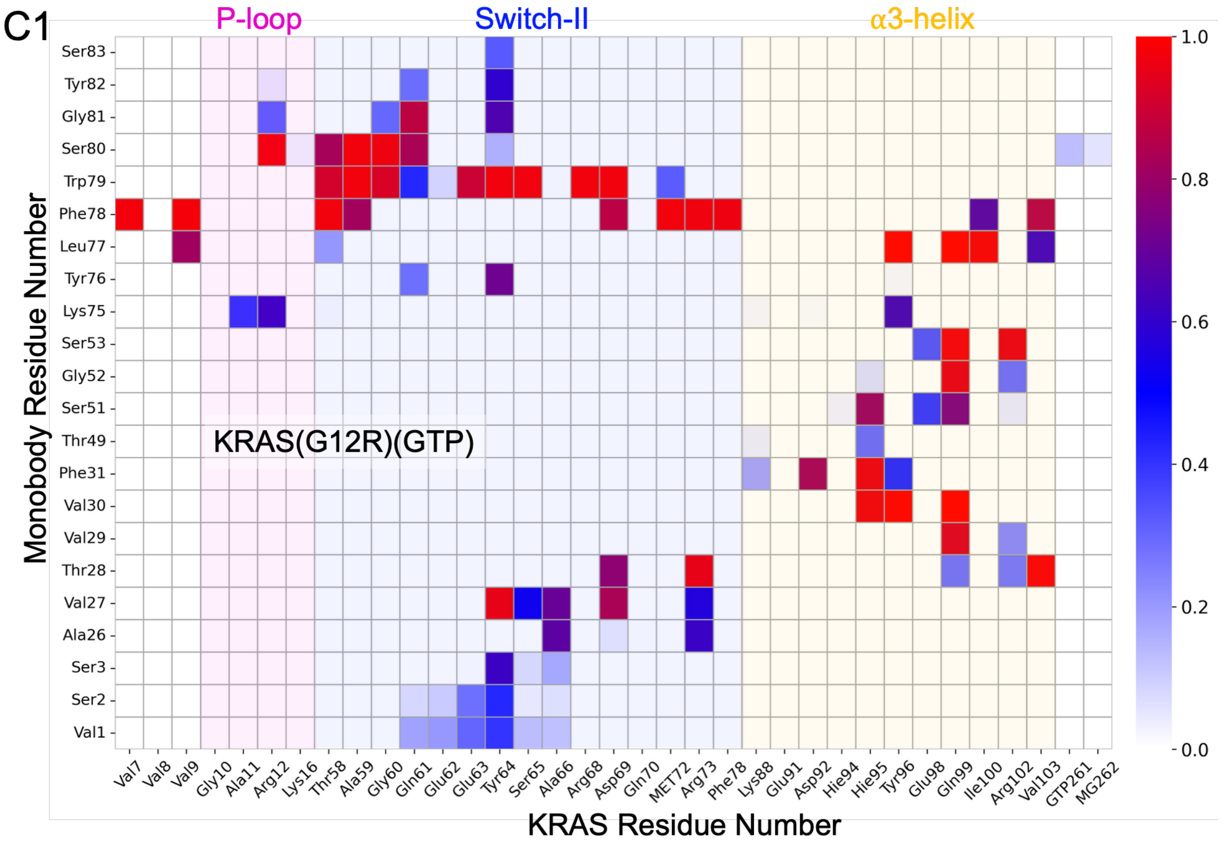
C1
P-loop
Switch-II
α3-helix
KRAS(G12R)(GTP)
Monobody Residue Number
KRAS Residue Number
Ser83
Tyr82
Gly81
Ser80
Trp79
Phe78
Leu77
Tyr76
Lys75
Ser53
Gly52
Ser51
Thr49
Phe31
Val30
Val29
Thr28
Val27
Ala26
Ser3
Ser2
Val1
Val7
Val8
Val9
Gly10
Ala11
Arg12
Lys16
Thr58
Ala59
Gly60
Gln61
Glu62
Glu63
Tyr64
Ser65
Ala66
Arg68
Asp69
Gln70
MET72
Arg73
Phe78
Lys88
Glu91
Asp92
Hie94
Hie95
Tyr96
Glu98
Gln99
Ile100
Arg102
Val103
GTP261
MG262
1.0
0.8
0.6
0.4
0.2
0.0

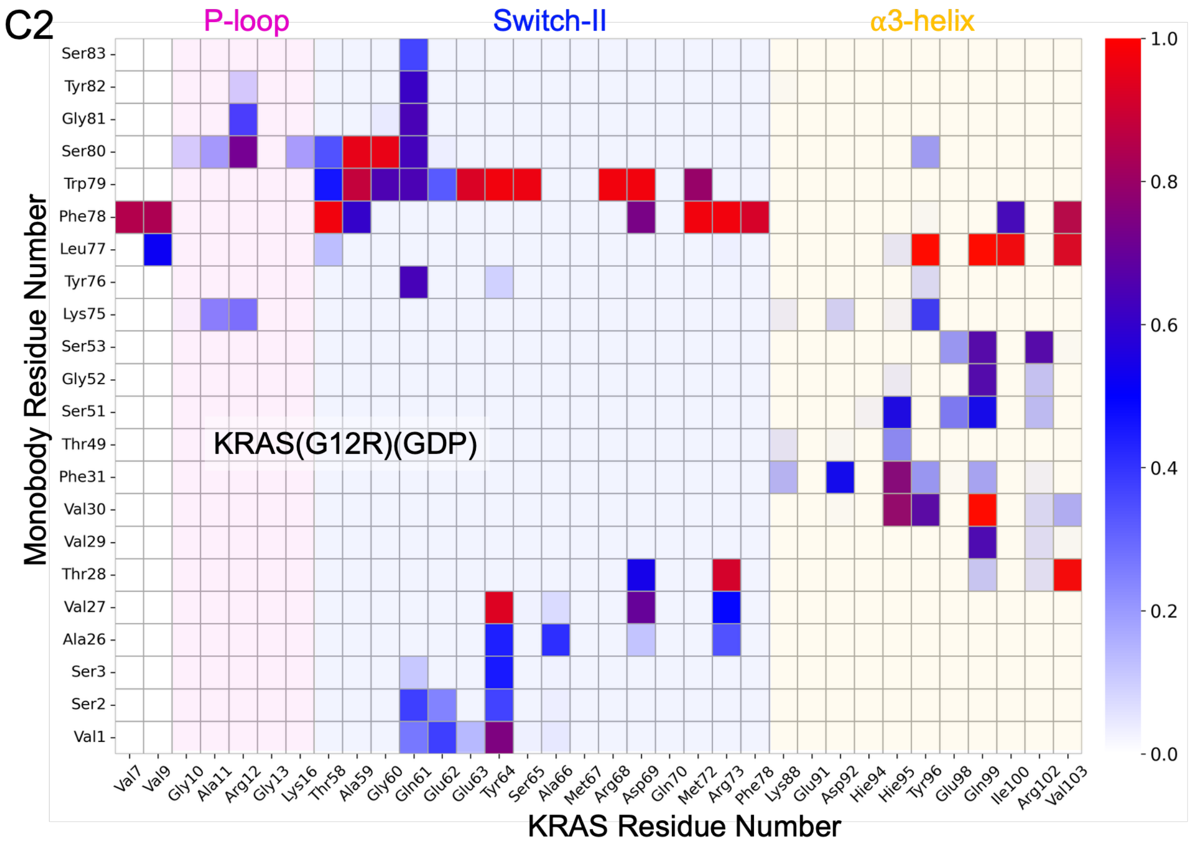
C2
P-loop
Switch-II
α3-helix
KRAS(G12R)(GDP)
Monobody Residue Number
KRAS Residue Number
Ser83
Tyr82
Gly81
Ser80
Trp79
Phe78
Leu77
Tyr76
Lys75
Ser53
Gly52
Ser51
Thr49
Phe31
Val30
Val29
Thr28
Val27
Ala26
Ser3
Ser2
Val1
Val7
Val9
Gly10
Ala11
Arg12
Gly13
Lys16
Thr58
Ala59
Gly60
Gln61
Glu62
Glu63
Tyr64
Ser65
Ala66
Met67
Arg68
Asp69
Gln70
Met72
Arg73
Phe78
Lys88
Glu91
Asp92
Hie94
Hie95
Tyr96
Glu98
Gln99
Ile100
Arg102
Val103
1.0
0.8
0.6
0.4
0.2
0.0

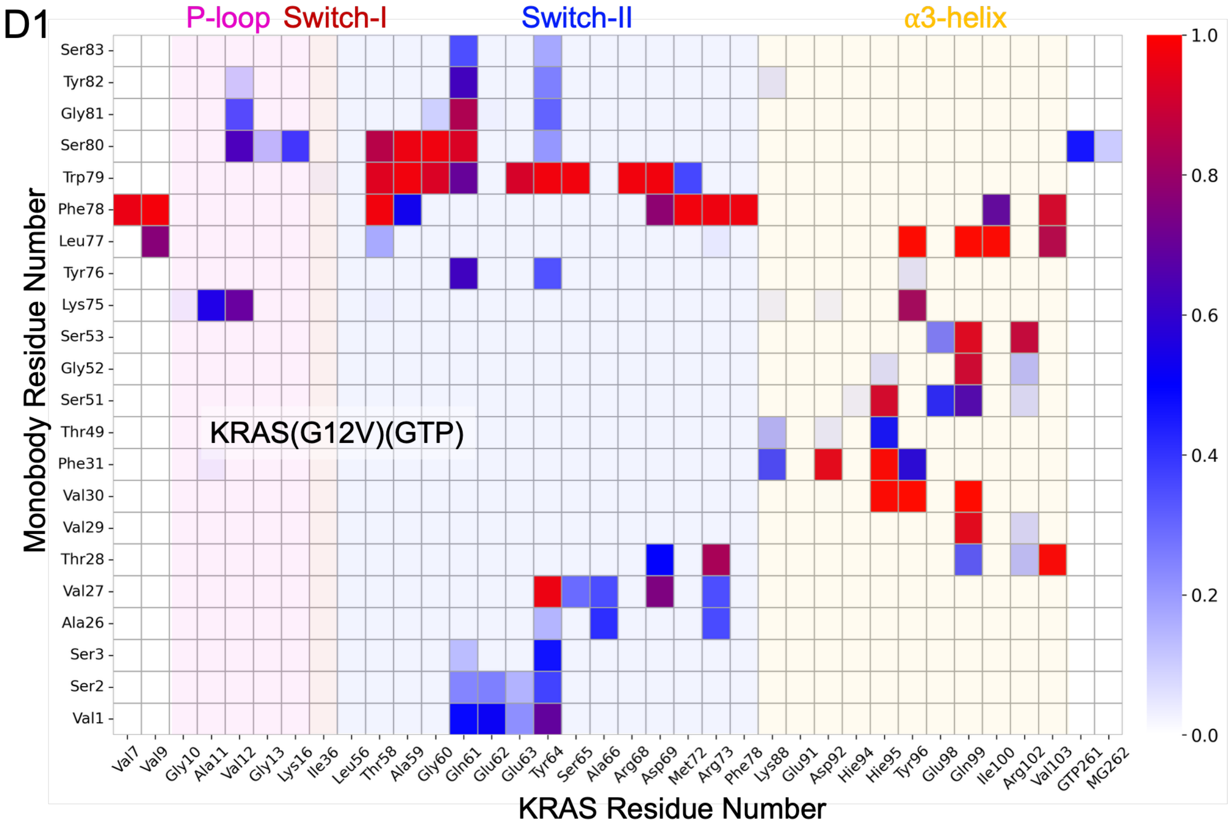
D1
P-loop
Switch-I
Switch-II
α3-helix
KRAS(G12V)(GTP)
Monobody Residue Number
Ser83
Tyr82
Gly81
Ser80
Trp79
Phe78
Leu77
Tyr76
Lys75
Ser53
Gly52
Ser51
Thr49
Phe31
Val30
Val29
Thr28
Val27
Ala26
Ser3
Ser2
Val1
Val7
Val9
Gly10
Ala11
Val12
Gly13
Lys16
Ile36
Leu56
Thr58
Ala59
Gly60
Gln61
Glu62
Glu63
Tyr64
Ser65
Ala66
Arg68
Asp69
Met72
Arg73
Phe78
Lys88
Glu91
Asp92
Hie94
Hie95
Tyr96
Glu98
Gln99
Ile100
Arg102
Val103
GTP261
MG262
KRAS Residue Number
1.0
0.8
0.6
0.4
0.2
0.0

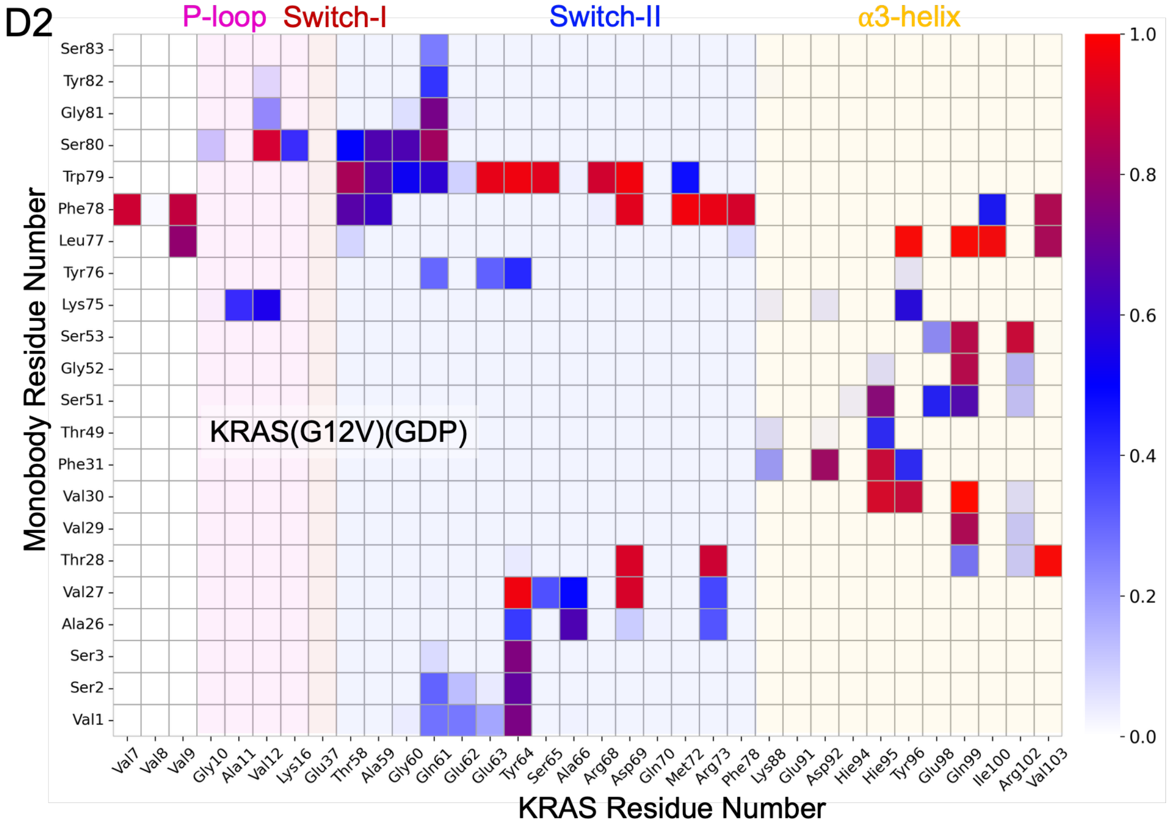
D2
P-loop
Switch-I
Switch-II
α3-helix
KRAS(G12V)(GDP)
Monobody Residue Number
Ser83
Tyr82
Gly81
Ser80
Trp79
Phe78
Leu77
Tyr76
Lys75
Ser53
Gly52
Ser51
Thr49
Phe31
Val30
Val29
Thr28
Val27
Ala26
Ser3
Ser2
Val1
Val7
Val8
Val9
Gly10
Ala11
Val12
Lys16
Glu37
Thr58
Ala59
Gly60
Gln61
Glu62
Glu63
Tyr64
Ser65
Ala66
Arg68
Asp69
Gln70
Met72
Arg73
Phe78
Lys88
Glu91
Asp92
Hie94
Hie95
Tyr96
Glu98
Gln99
Ile100
Arg102
Val103
KRAS Residue Number
1.0
0.8
0.6
0.4
0.2
0.0

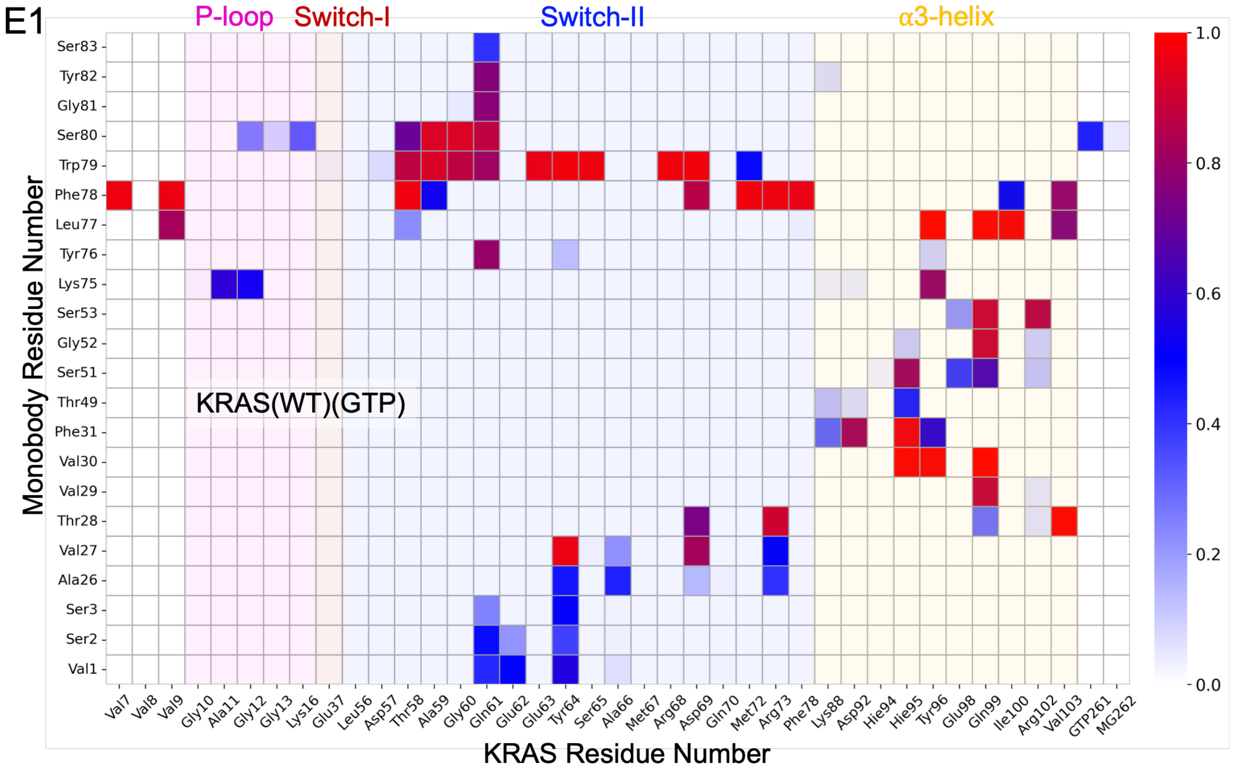
E1
P-loop
Switch-I
Switch-II
α3-helix
KRAS(WT)(GTP)
Monobody Residue Number
KRAS Residue Number
Ser83
Tyr82
Gly81
Ser80
Trp79
Phe78
Leu77
Tyr76
Lys75
Ser53
Gly52
Ser51
Thr49
Phe31
Val30
Val29
Thr28
Val27
Ala26
Ser3
Ser2
Val1
Val7
Val8
Val9
Gly10
Ala11
Gly12
Gly13
Lys16
Glu37
Leu56
Asp57
Thr58
Ala59
Gly60
Gln61
Glu62
Glu63
Tyr64
Ser65
Ala66
Met67
Arg68
Asp69
Gln70
Met72
Arg73
Phe78
Lys88
Asp92
Hie94
Hie95
Tyr96
Glu98
Gln99
Ile100
Arg102
Val103
GTP261
MG262
1.0
0.8
0.6
0.4
0.2
0.0

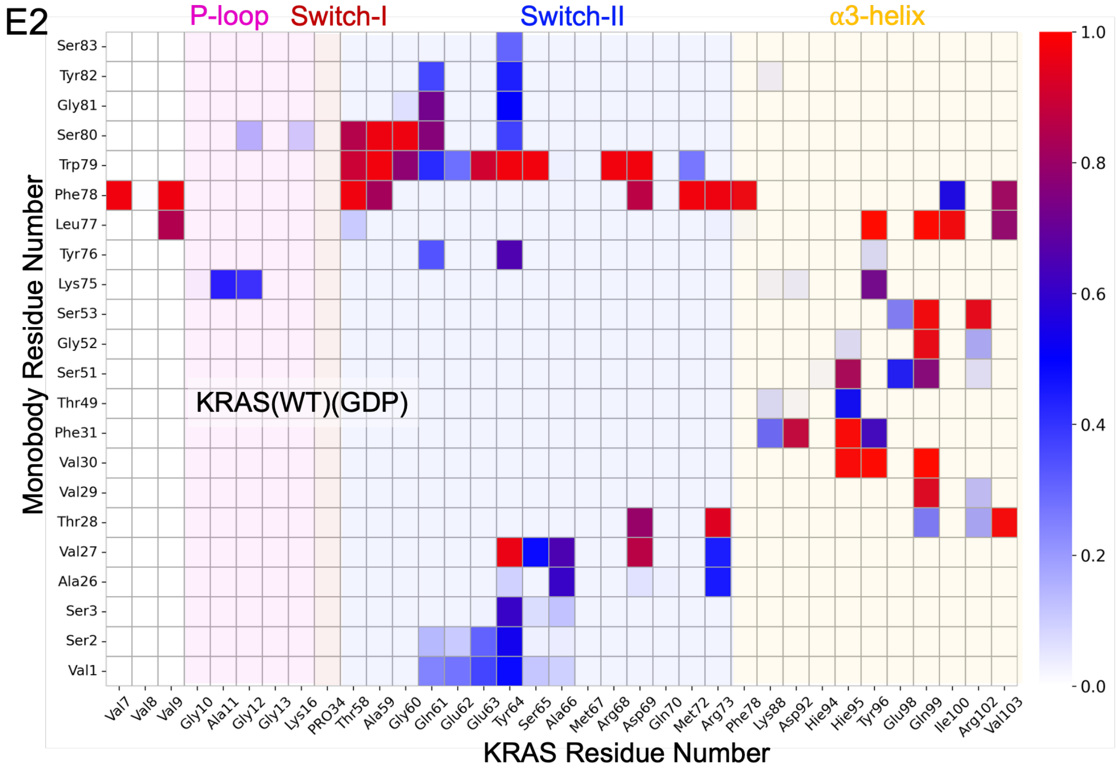
E2
P-loop
Switch-I
Switch-II
α3-helix
KRAS(WT)(GDP)
Monobody Residue Number
KRAS Residue Number
Ser83
Tyr82
Gly81
Ser80
Trp79
Phe78
Leu77
Tyr76
Lys75
Ser53
Gly52
Ser51
Thr49
Phe31
Val30
Val29
Thr28
Val27
Ala26
Ser3
Ser2
Val1
Val7
Val8
Val9
Gly10
Ala11
Gly12
Gly13
Lys16
PRO34
Thr58
Ala59
Gly60
Gln61
Glu62
Glu63
Tyr64
Ser65
Ala66
Met67
Arg68
Asp69
Gln70
Met72
Arg73
Phe78
Lys88
Asp92
Hie94
Hie95
Tyr96
Glu98
Gln99
Ile100
Arg102
Val103
1.0
0.8
0.6
0.4
0.2
0.0

**Figure S6:** Percentage contact occupancy for residue pairs at the KRAS-12D4 interface. Panels A1 and A2 correspond to KRAS(G12D) in the GTP- and GDP-bound states, respectively; panels B1 and B2 correspond to KRAS(G12C); panels C1 and C2 correspond to KRAS(G12R); panels D1 and D2 correspond to KRAS(G12V); and panels E1 and E2 correspond to KRAS(WT), each shown in the GTP- and GDP-bound states. Major KRAS structural regions are color-coded to highlight their contributions to the binding interface. This figure is supplemental to Figure 6.

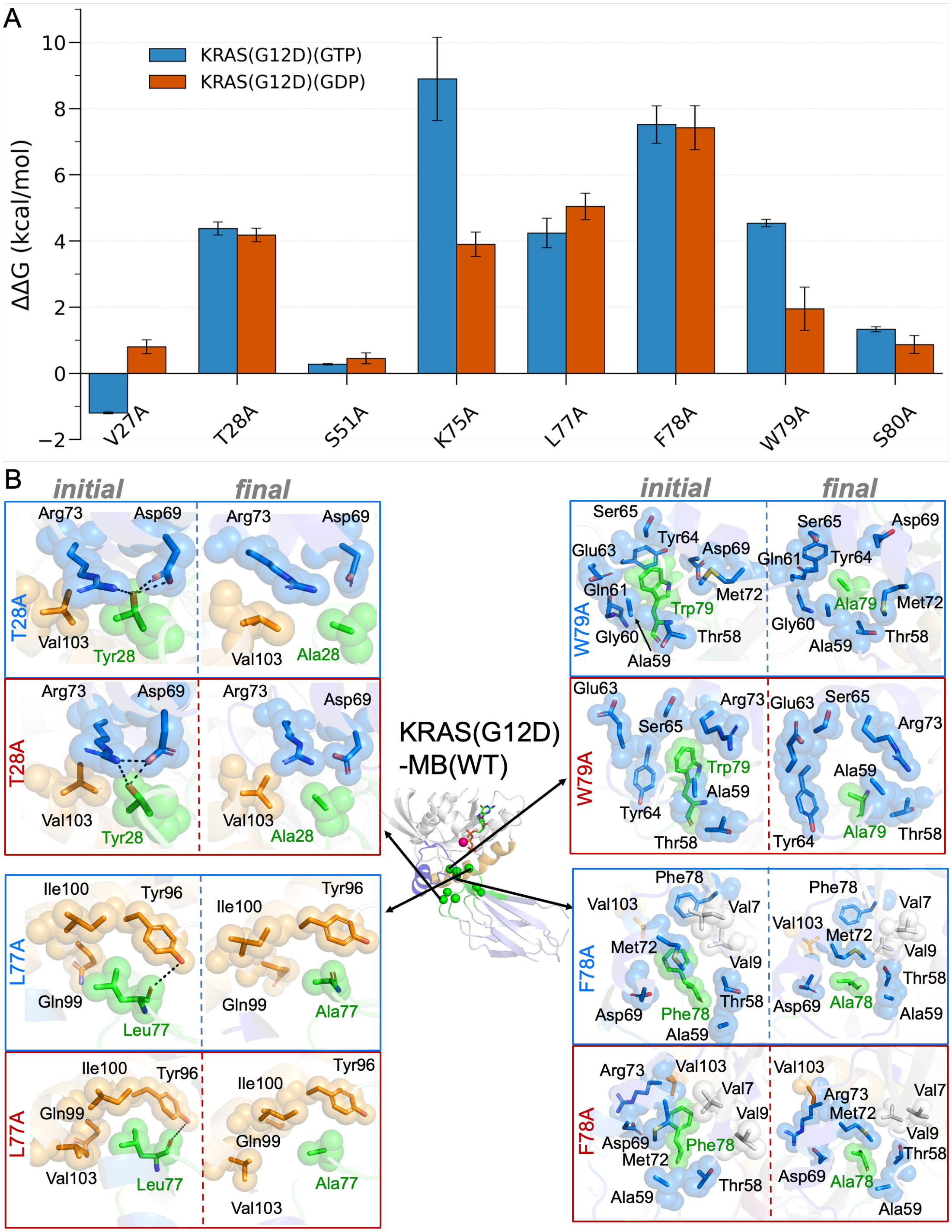


**Figure S7:** Energetic effects of alanine mutations in the monobody and associated structural changes at the KRAS-monobody interface. (A) Relative binding free-energy changes (ΔΔG) for alanine mutations of key monobody interface residues interacting with KRAS(G12D) in the GTP- and GDP-bound states. Positive ΔΔG values indicate reduced binding affinity relative to the parental WT 12D4 monobody. Error bars represent standard error of mean obtained from 3 independent calculations. (B) Structural comparison of different 12D4 mutants. Initial and final representative structures from MD simulations are shown for T28A, L77A, F78A, and W79A monobody mutations. Monobody residues are shown in green sticks. These mutations disrupt local interactions at the KRAS-monobody interface and reduce the monobody binding energy relative to the WT monobody.

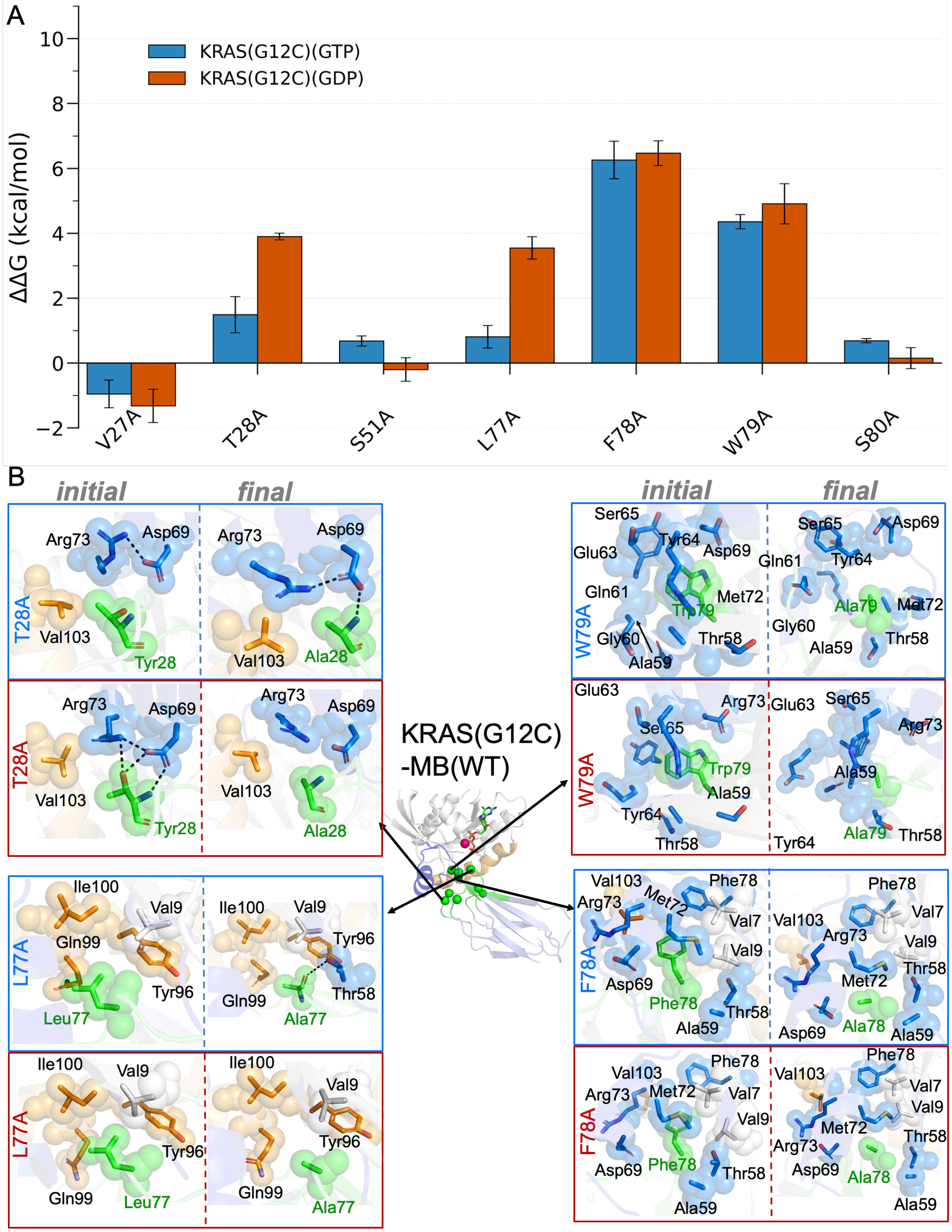


**Figure S8:** Data similar to Figure S7 are shown for alanine mutations in the monobody (A) and associated structural changes at the KRAS(G12C)-monobody interface (B).

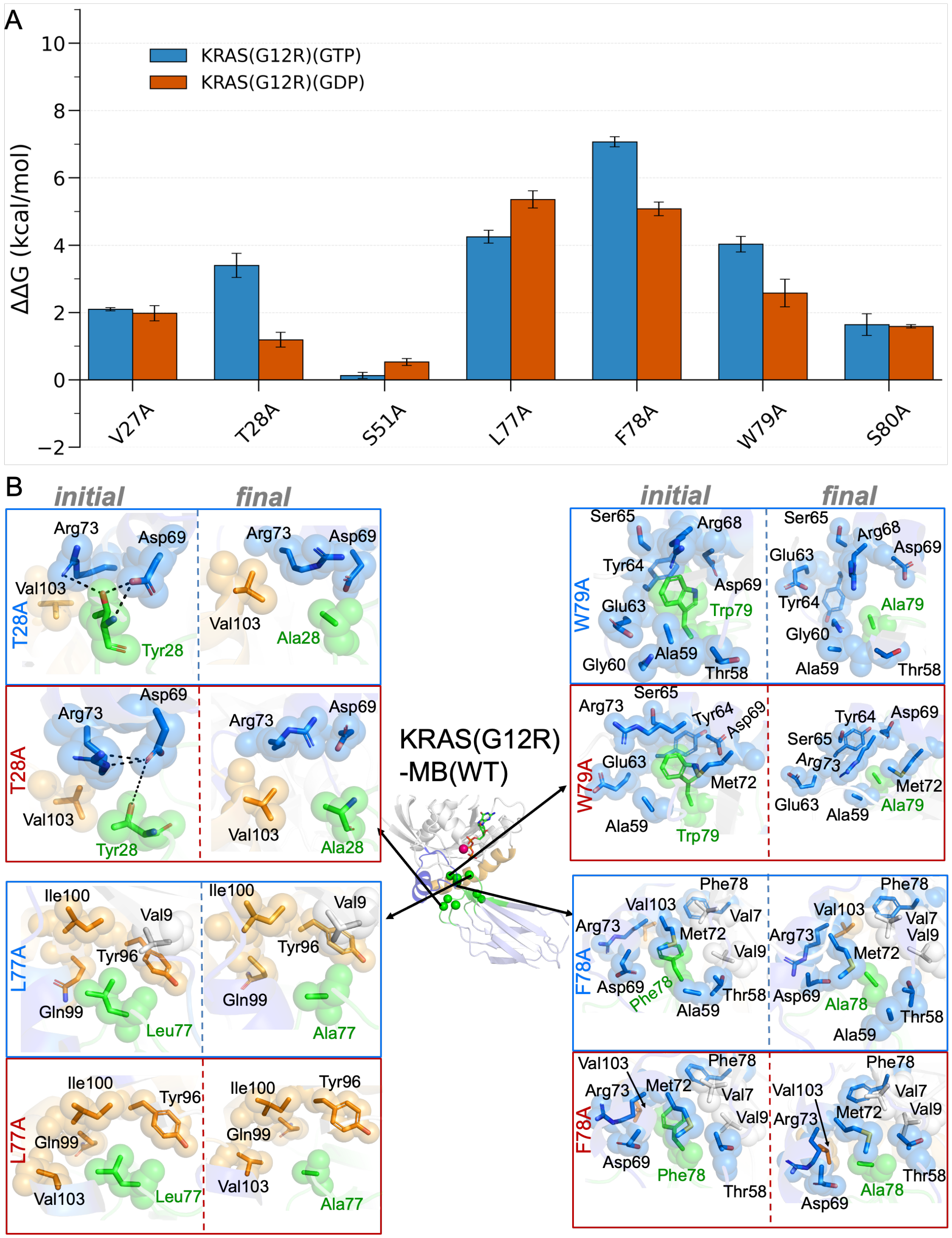


**Figure S9:** Data similar to Figure S7 are shown for alanine mutations in the monobody (A) and associated structural changes at the KRAS(G12R)-monobody interface (B).

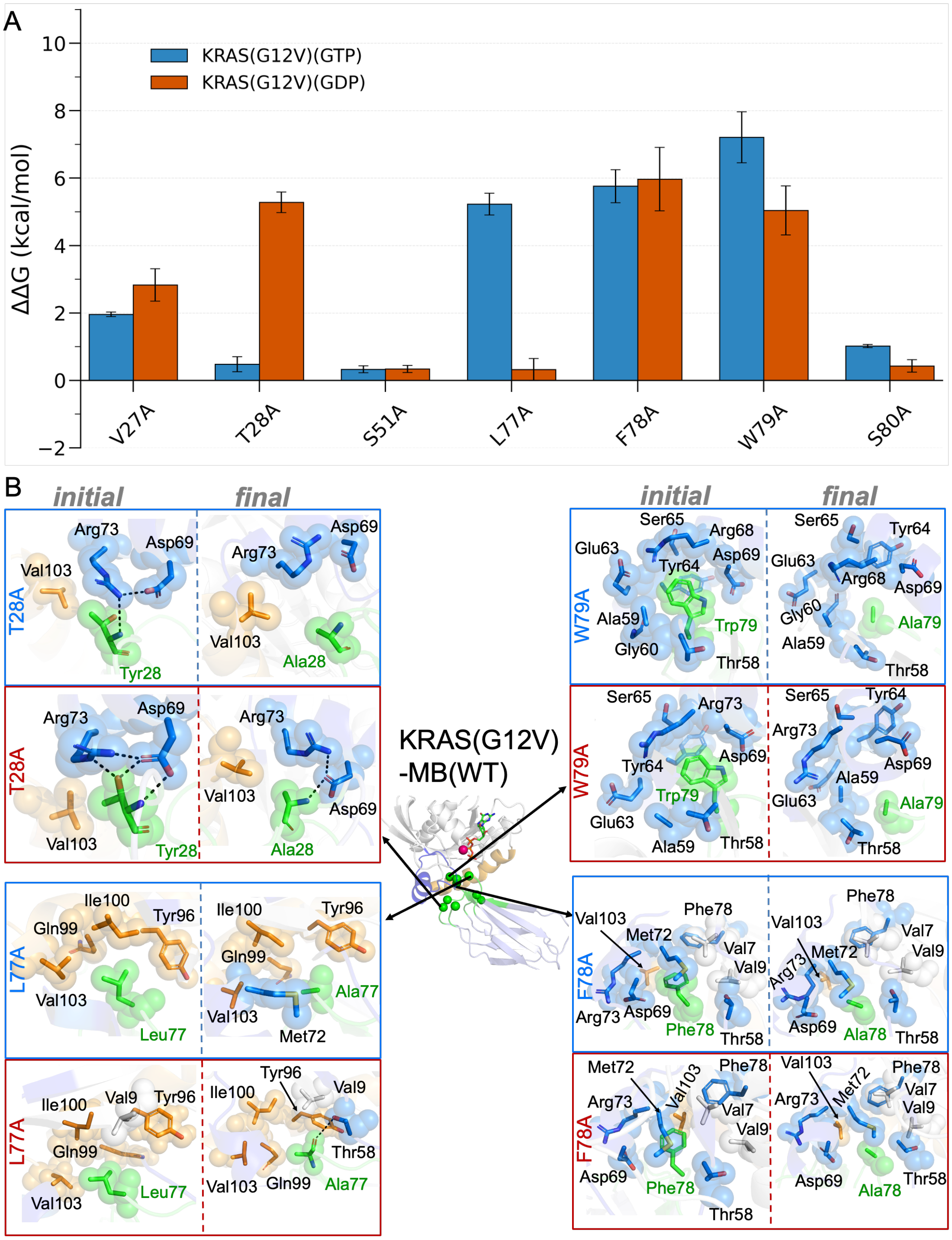


**Figure S10:** Data similar to Figure S7 are shown for alanine mutations in the monobody (A) and associated structural changes at the KRAS(G12V)-monobody interface (B).

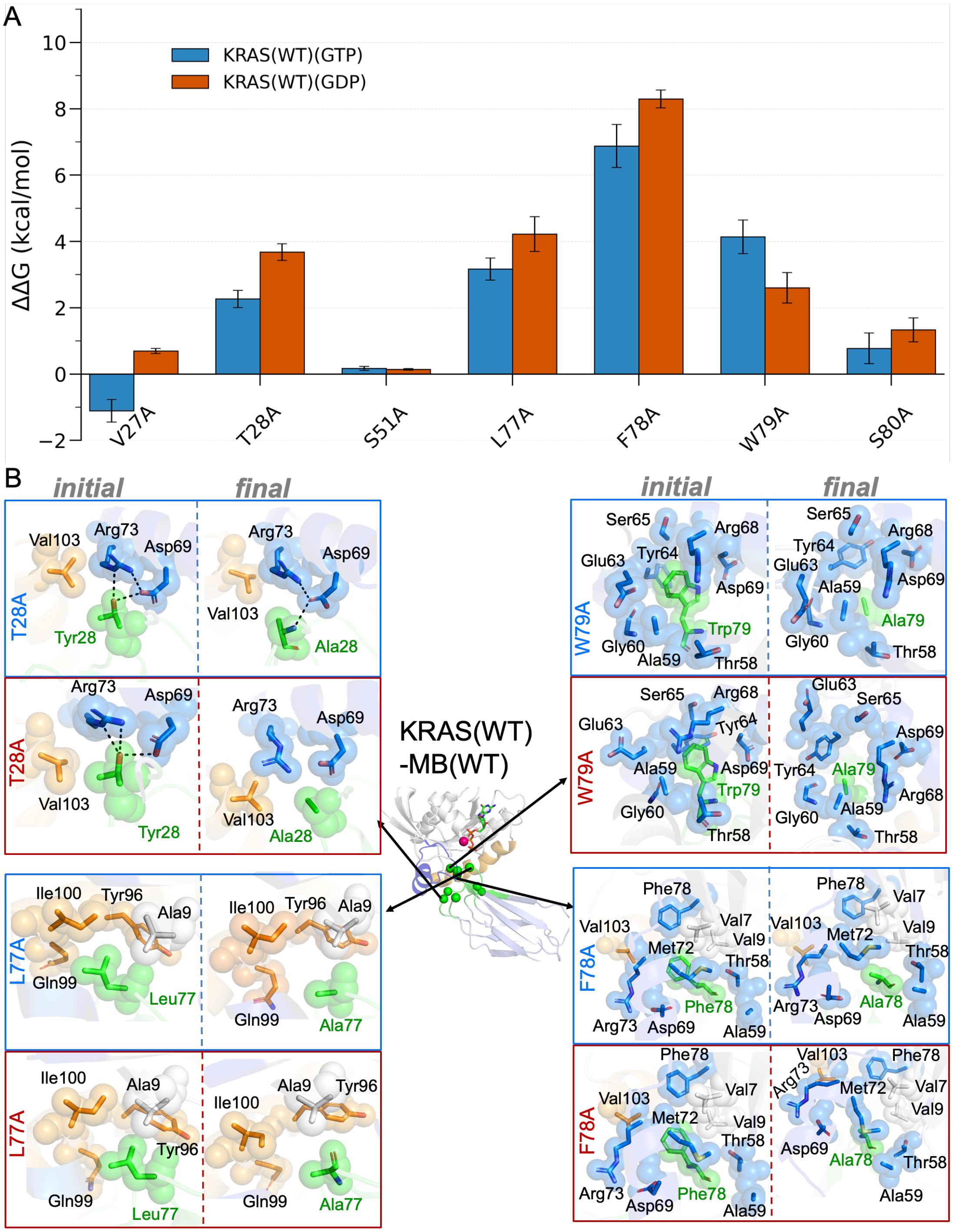


**Figure S11:** Data similar to Figure S7 are shown for alanine mutations in the monobody (A) and associated structural changes at the KRAS(WT)-monobody interface (B).
